# Deciphering the fibre-orientation independent component of R_2_* (R_2,iso_*) in the human brain with a single multi-echo gradient-recalled-echo measurement under varying microstructural conditions

**DOI:** 10.1101/2022.03.28.486076

**Authors:** Francisco J. Fritz, Laurin Mordhorst, Mohammad Ashtarayeh, Joao Periquito, Andreas Pohlmann, Markus Morawski, Carsten Jaeger, Thoralf Niendorf, Kerrin J. Pine, Martina F. Callaghan, Nikolaus Weiskopf, Siawoosh Mohammadi

## Abstract

The effective transverse relaxation rate (R_2_*) is sensitive to the microstructure of the human brain, e.g. the g-ratio characterising the relative myelination of axons. However, R_2_* depends on the orientation of the fibres relative to the main magnetic field degrading its reproducibility and that of any microstructural derivative measure. To decipher its orientation-independent part (R_2,iso_*), a second-order polynomial in time (M2) can be applied to single multi-echo gradient-recalled-echo (meGRE) measurements at arbitrary orientation. The linear-time dependent parameter, *β*_1_, of M2 can be biophysically related to R_2,iso_* when neglecting the signal from the myelin water (MW) in the hollow cylinder fibre model (HCFM). Here, we examined the effectiveness of M2 using experimental and simulated data with variable g-ratio and fibre dispersion. We showed that the fitted *β*_1_ effectively estimates R_2,iso_*when using meGRE with long maximum echo time (TE_max_ ≈ 54 ms) but its microscopic dependence on the g-ratio was not accurately captured. This error was reduced to less than 12% when accounting for the MW contribution in a newly introduced biophysical expression for *β*_1_. We further used this new expression to estimate the MW fraction (0.14) and g-ratio (0.79) in a human optic chiasm. However, the proposed method failed to estimate R_2,iso_* for a typical *in-vivo* meGRE protocol (TE_max_ ≈ 18 ms). At this TE_max_ and around the magic angle, the HCFM-based simulations failed to explain the R_2_*-orientation-dependence. In conclusion, estimation of R_2,iso_* with M2 *in vivo* requires meGRE protocols with very long TE_max_ ≈ 54 ms.

## 1. Introduction

The effective transverse relaxation rate (R_2_* = 1/T_2_*) is a nuclear magnetic resonance (NMR) relaxation property (Tofts, 2004) that enables non-invasive characterisation of the microstructure of the human brain (Does, 2018; MacKay et al., 2006; Weiskopf et al., 2021). The microstructural sensitivity of R_2_* makes it particularly interesting for neuroscience and clinical research studies (Callaghan et al., 2014; Draganski et al., 2011; Kirilina et al., 2020; Langkammer et al., 2010). This is because R_2_* is sensitive not only to free and myelin water pools in the brain (Dula et al., 2010a; MacKay et al., 2006; Weiskopf et al., 2021) but also to microscopic perturbations in the main magnetic field 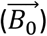 (Chavhan et al., 2009). These microscopic perturbations are caused by the different magnetic susceptibilities of biological structures (Duyn and Schenck, 2017) like the diamagnetic myelin sheath (Alonso-Ortiz et al., 2018; Duyn, 2014; Kucharczyk et al., 1994; Lee et al., 2017) and paramagnetic iron deposits in glial cells (Li et al., 2009; Ordidge et al., 1994; Yao et al., 2009). Moreover, it has been shown that R_2_* is also strongly dependent on the angular orientation of the white matter fibre tracts relative to 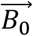 (Lee et al., 2011,2012) confounding the mapping of R_2_* to the underlying microstructure. The angular orientation dependence of R_2_* can be decomposed into an isotropic, i.e. angular-independent, component (R_2,iso_*) and an angular-dependent component using either gradient-recalled echo (GRE) acquisitions at several angular orientations (Oh et al., 2013; Rudko et al., 2014; Wharton and Bowtell, 2013) or hybrid diffusion weighted imaging (DWI) and GRE acquisitions with reduced numbers of distinct angular-orientations (Gil et al., 2016). However, both methods are impractical for clinical research due to the constrained and inconvenient positioning of the patient’s head in the radiofrequency receiver coil needed to achieve the required distinct angular orientations.

Recently, it was shown that R_2,iso_*, with interpretable microstructural information of the myelinated fibres, can be obtained from a single multi-echo GRE (meGRE) measurement (Papazoglou et al., 2019). In this work, they used the hollow cylinder fibre model (HCFM, (Wharton and Bowtell, 2013, 2012)) to derive a second-order approximation of the logarithm of the time-dependent signal where the linear component in time (*β*_1_) was a proxy for R_2,iso_* and the orientation-dependent part was regressed out by the second-order term in time (*β*_2_). In the following, this model is denoted as the log-quadratic model (M2) to be distinguished from the classical log-linear model (M1, (Elster, 1993; Peters et al., 2007; Weiskopf et al., 2014)), where the linear parameter in time (*α*_1_) contains contributions from both R_2,iso_* and the angular-dependent part of R_2_*.

Since the M2-proxy for R_2,iso_* (i.e. the *β*_1_ parameter) is based on the HCFM, it can be directly related to microscopic tissue properties. In the HCFM model, the dephasing is caused by the hollow-cylinder fibre and is mainly driven by its g-ratio, which is defined as the ratio between the inner and the outer radii of a myelinated axon (Rushton, 1951). In this model, all potential other perturbers are ignored (e.g. non-local effects of susceptibility inhomogeneities due to cavities, vessels, and iron molecules). Thus, the predicted *β*_1_ parameter depends only on the transverse relaxation rate of the free water molecules of the non-myelinated compartments (R_2N_), i.e. inside (intra-) and outside (extra-) of the axonal cell. A counter-intuitive prediction of M2 is that its *β*_1_ parameter is independent of any changes associated with the myelin water signal, e.g., changes in the myelin water fraction (MWF). This independence of the *β*_1_ parameter to MWF could contradict the hypothesis that R_2,iso_* can be biologically modelled via *β*_1_, since R_2,iso_* has been shown to be linearly dependent on MWF, see (Kirilina et al., 2020; Lee et al., 2017). Moreover, M2 assumes that axonal fibres are perfectly aligned or even described by one representative axon. However, most of the fibre bundles in the human brain possess a diverse range of topographies, i.e. show fanning and bending, or mildly to acute crossing, e.g. (Jeurissen et al., 2019; Schmahmann et al., 2009, 2007) and different levels of relative myelination, e.g. (Mohammadi et al., 2015). Besides that, the performance of M2 in estimating R_2,iso_* via *β*_1_ has only been tested with data incorporating very long maximum echo times of ≈ 54 ms (Papazoglou et al., 2019). Such a long maximum echo time, is unusual for *in vivo* meGRE measurements with whole-brain coverage (Weiskopf et al., 2013; Ziegler et al., 2019) because it increases the total scan time as well as the propensity for bulk and physiological motion.

This work explores the potential and pitfalls of using M2 to estimate R_2,iso_* via *β*_1_, from a single-orientation meGRE, while varying biological fibre properties and maximum echo times. Moreover, it tests the counter-intuitive hypothesis, based on M2, that the estimated *β*_1_ is independent of the MWF. To this end, we use simulated (hereafter *in silico*) data and *ex vivo* MRI. The *in silico* data were simulated using the HCFM to generate realistic meGRE datasets from an ensemble of myelinated axons, for which the ground truth biophysical parameters (i.e., g-ratio, fibre dispersion and angular orientation) are known and can be varied. The *ex vivo* dataset combines high-resolution DWI and multi-orientation meGRE imaging of a human optic chiasm to generate gold-standard datasets where the fibre orientation and dispersion are known. Both datasets are used to perform the following analyses: First, we assess the capability of M2 to estimate R_2,iso_* via *β*_1_ for varying g-ratio values and fibre dispersions. Second, we assess the microstructural interpretability of *β*_1_. To this end, we test the hypothesis that *β*_1_ is independent of MWF by evaluating the deviation between fitted *β*_1_ using the *in silico* data and the biophysically-predicted *β*_1_ by M2. Additionally, we perform the same comparison as above using a novel heuristic expression that incorporates the MWF dependence into the predicted *β*_1_. Third, we demonstrate that the heuristic expression for *β*_1_ can be used to calculate MWF and the g-ratio from the *β*_1_ of the *ex vivo* data. And, fourth, we assess the capability of M2 to estimate R_2,iso_* via *β*_1_ for shorter maximal echo times more typical of *in vivo* meGRE applications.

## 2. Background

### 2.1 Overview of the hollow cylinder fibre model and the approximated log -quadratic model

The hollow cylinder fibre model (HCFM, (Wharton and Bowtell, 2013, 2012)) proposes an analytical approximation describing the dependence of the GRE signal on the angular orientation 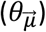 defined as the angle between the main magnetic field 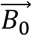 and the hollow-cylinder fibre 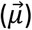. This approximation establishes that the total MR signal comes from water molecules in an *infinitely long* hollow cylinder affected by the diamagnetic myelin sheath (Liu, 2010). The diamagnetic myelin sheath magnetically perturbs the water molecules in three distinct compartments: (1) the intra-axonal (S_A_), (2) myelin (S_M_) and (3) extra-cellular (S_E_) compartments (details in Appendix, section 9.1). When the signal of the water molecules in the myelin compartment is neglected, the signal magnitude of the HFCM can be approximated by a log-quadratic model (M2) in time (Papazoglou et al., 2019):

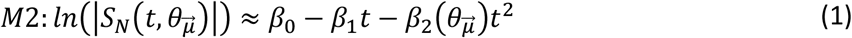

where *β*_0,1,2_ are the model parameters and S_N_ is the non-myelin signal (i.e., S_N_ = S_A_ + S_E_). In this model, the slope *β*_1_ is considered as a proxy for R_2,iso_* because it does not possess any 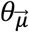 dependence (Equations A16b), whereas *β*_2_ contains all the 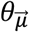 dependent information of R_2_* (Equations A16c, details in Appendix, section 9.3).

In contrast to M2, the slope (*α*_1_) in the classic log-linear model (M1, (Elster, 1993)) of R_2_* is a function of R_2,iso_* and the 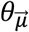 dependent components of R_2_* (e.g. see (Lee et al., 2012b, 2011)):

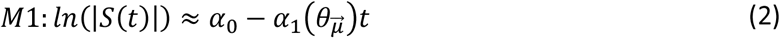

### 2.2 Myelin dependence of *β*_1_ parameter as predicted by the log-quadratic model (M2)

The slope *β*_1_ of M2, which is considered to be a proxy for R_2,iso_*, is derived from the HCFM of a two-pool system in the slow-exchange regime: a fast decaying water pool consisting of the myelin water with a relaxation rate *R*_2*M*_ and a slow decaying water pool with a relaxation rate *R*_2*N*_ consisting on the intra and extra cellular water. In this model, the only source of dephasing is caused by the hollow-cylinder fibre and all potential other perturbers are ignored (e.g. non-local effects of susceptibility inhomogeneities due to cavities, vessels, and iron molecules). Consequently in the approximation of M2 (Equation A16b, section 9.3), the predicted *β*_1_ parameter (hereafter *β*_1,*nm*_) is given by the transverse relaxation rate of the non-myelin water pool (*R*_2*N*_):

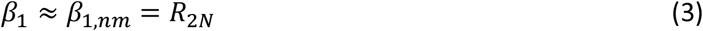

We hypothesise here that for realistic tissue composition (i.e. g-ratio equal to or smaller than 0.8), where the myelin compartment cannot be neglected, Equation 3 is invalid. This hypothesis is supported by previous observations showing that R_2,iso_* depends on the myelin water fraction, MWF (e.g.(Lee et al., 2017; Weber et al., 2020)).

Here, we propose an alternative heuristic biophysical expression of the predicted *β*_1_ parameter (hereafter *β*_1,*m*_), also based on the HCFM (Equation A17) but without assuming the myelin compartment to be negligible. In this case, the expected dependence of R_2,iso_* on variation in the MWF remains:

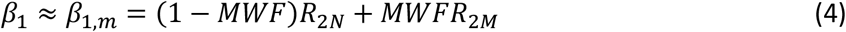

where *R*_2*M*_ is the relaxation rate of the myelin water pool. It follows from the heuristic model that the fitted *β*_1_ is a weighted sum of the relaxation rates of the two pools, and that Equation 3 (*β*_1,*nm*_) is a special case of Equation 4 (*β*_1,*m*_) when the MWF is equal to 0.

Based on our hypothesis, we expect that the heuristic expression for *β*_1,*m*_ can better describe the fitted *β*_1_ when varying the g-ratio, and thus is a better proxy of R_2,iso_*.

Finally, we describe the dependence of MWF on the g-ratio by redefining the compartmental volumes: intra-axonal V_A_, extra-axonal V_E_ and myelin V_M_ (Equation 18a), as a function of the g_ratio_ and fibre volume fraction (FVF) as: V_A_ = FVF · g^2^_ratio_, V_E_ = 1 – FVF, and V_M_ = FVF · (1 – g^2^_ratio_). If the proton densities of the non-myelinated compartments are equal (ρ_A_ = ρ_E_ = ρ_N_), then the MWF can be rewritten as:

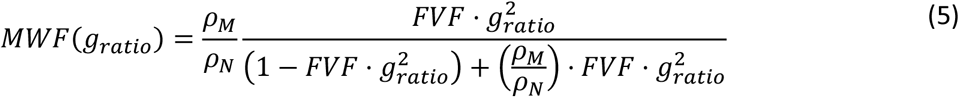

Therefore, the g-ratio could be estimated from the MWF if the proton densities and FVF were known.

## 3. Methods

This section explains the approaches used for data acquisition, data analysis and for comparing the results obtained from the *ex vivo* data and the findings derived from the *in silico* data.

### 3.1. Ex-vivo: Optic chiasm

#### 3.1.1. Sample and data acquisition

A human optic chiasm (OC) from a patient without any diagnosed neurological disorder was measured (male, 59 years, multi-organ failure, 48 hours *postmortem* interval, ∼80 days of fixation in phosphate buffered saline (PBS) pH 7.4 with 0.1% sodium acide NaN3 containing 3% PFA + 1% GA) with prior informed consent (Ethical approval #205/17-ek). Two MR techniques were used: R_2_*-weighted GRE and diffusion-weighted MRI (dMRI).

All R_2_*-weighted GRE acquisitions were performed on a 7 T Siemens Magnetom MRI scanner (Siemens Healthcare GmbH, Erlangen, Germany) using a custom 2-channel transmit/receive circularly polarized (CP) coil with a diameter of 60 mm. The OC sample was fixed within an acrylic sphere of 60 mm diameter filled with agarose (1.5% Biozym Plaque low melting Agarose, Merck, Germany) dissolved in PBS (pH 7.4 + 0.1% sodium) and scanned at sixteen orientations (covering a solid angle, with azimuthal and elevation angles from 0° to 90°, Figure 1A) using the 3D multi-echo GRE (meGRE) MRI (hereafter: **GRE dataset**). For each angular meGRE measurement, sixteen echoes were acquired at equally spaced echo times (TE) ranging from 3.4 to 53.5 ms (increment 3.34 ms) with a repetition time (TR) of 100 ms, a field of view (FoV) of (39.00 mm)^3^, a matrix size of 112^3^, resulting in an isotropic voxel resolution of (0.35 mm)^3^, non-selective RF excitation with a flip angle of 23° and a gradient readout bandwidth of 343 Hz/px. The acquired MR data are the same as reported in (Papazoglou et al., 2019).

**Figure 1:**
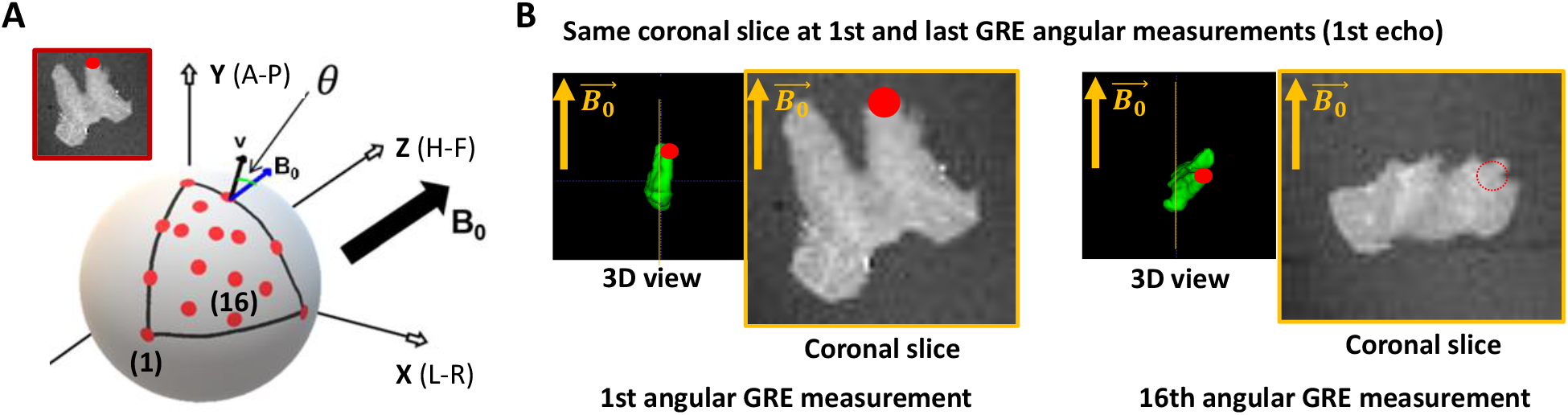
Acquisition of the multi-angular multi-echo gradient recalled echo (meGRE) ex vivo data. (A) An illustration of the different angular measurements performed on the optic chiasm (OC) specimen. The red dots show the position of the optical tracts (see inset) for the different measurements. The different coordinates (spatial, x-y-z and anatomical, anterior-head-right, A-H-R) are shown. (B) Illustration of the first echo meGRE image acquired at the first and last angular measurement. The 3D view shows the specimen position to the main magnetic field 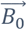 and the position of the optical tract (red dot). The yellow line shows the same coronal slice image.

Additionally, multi-shell dMRI data (hereafter: **dMRI dataset**), suitable for NODDI analysis, were acquired on a 9.4 T small animal MR system (Bruker Biospec 94/20; Bruker Biospin, Ettlingen, Germany) using a 2-channel receiver cryogenically cooled quadrature transceiver surface RF coil (Bruker Biospin, Ettlingen, Germany) and a gradient system with B_max_ = 700 mT/m per gradient axis. This dataset was acquired with a slice-selective (2D) pulsed-gradient spin-echo (PGSE) technique, consisting of four diffusion-weighting shells (number of directions) of b = 1000 s/mm^2^ (60), 4000 s/mm^2^ (60), 8000 s/mm^2^ (60) and 12000 s/mm^2^ (60) with 35 non-diffusion-weighted volumes (∼ 0 s/mm^2^). The fixed diffusion parameters were diffusion time Δ = 13 ms, diffusion gradient duration δ = 6 ms. The remaining sequence parameters were TE = 27 ms, TR = 30 s (to acquire all the slices), FoV = 20.75 × 16.00 × 12.50 mm^3^, matrix size = 83 × 64 × 50, isotropic voxel resolution = (0.25 mm)^3^, slice selective pulses with flip angles of 90° (excitation) and 180° (refocusing) and receiver bandwidth of 9411 Hz/px.

#### 3.1.2. Dispersion and mean fibre orientation estimation from dMRI dataset

To incorporate the voxel-wise information regarding the angular orientation of the fibres to 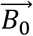 and fibre’s dispersion, the dMRI datasets were analysed with two diffusion models: Neurite Orientation Dispersion and Density Imaging (NODDI) (Zhang et al., 2012) and Diffusion Tensor Imaging (DTI) (Basser et al., 1994). The NODDI toolbox was adjusted for *ex vivo* analysis (Wang et al., 2019) and used all the diffusion shells. The main neurite (hereafter fibre) orientation 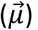, a measure of the fibre dispersion (κ), and fibre density (volume fraction of the intracellular compartment, ICVF) maps were estimated from this analysis. The DTI model used the first two diffusion shells (b-values: 1000 s/mm^2^ and 4000 s/mm^2^) and only the fractional anisotropy (FA) map was estimated, because this map was used only for diffusion-to-GRE coregistration (section 3.1.3).

#### 3.1.3 Coregistration of the GRE angular measurements and dMRI results

To establish a voxel-to-voxel relationship between the meGRE signal at different angular orientations and the properties estimated from dMRI, i.e., κ, 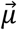 and ICVF, we coregistered the angular meGRE measurements and the dMRI measurement. To this end, we estimated two sets of transformation matrices: first, transformation matrices that coregister the angular measurements in GRE space (see Figure 1A); and second, a transformation matrix that coregister from GRE space to dMRI space (see Figure 1B). The coordinate system of GRE space was defined by the first meGRE angular measurement.

To estimate the transformation matrices that coregister the angular meGRE measurements to the first angular meGRE measurement, a manual coregistration was performed and refined later with an automatic coregistration. This pair of coregistrations resulted in the transformation matrix *T*_*GRE*:*i*,1_ (i = 2 … 16). When aligning the meGRE volumes, the respective 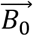 directions had to be adjusted accordingly. This was done by aligning the 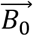 direction of the *i*-th meGRE using the respective transformation matrix *T*_*GRE*:*i*,1_ (see insets). Both coregistrations were performed using the 3D Slicer software (http://www.slicer.org and (Fedorov et al., 2012)).

To estimate the transformation matrix from GRE to dMRI space, the FA map from the DTI analysis (section 3.1.2) was coregistered to the first meGRE angular measurement (Figure 1B). This transformation matrix, *T*_*Diff,GRE*_, was applied to coregister the NODDI results, i.e., κ, 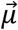 and ICVF, to the GRE space. However, this transformation matrix was also used to align the sixteen new 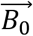 estimated from the angular meGRE coregistration to the dMRI space (section 3.1.4). This GRE-to-diffusion transformation was performed using the coregistration module in SPM 12 (http://www.fil.ion.ucl.ac.uk/spm).

#### 3.1.4. Voxel-wise estimation of the angular orientation, 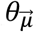, between fibres and 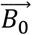

The angular orientation 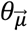 between fibres and 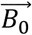 for each meGRE angular measurement was calculated in dMRI space by computing the arccosine of the inner product between 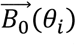 and 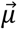, i.e., 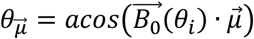 (Figure 3C). In this computation, 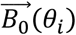 is the resulting 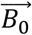 after the transformation from the i-th meGRE angular measurement to the first meGRE angular measurement (*T*_*GRE*:*i*,1_), and the transformation from GRE to dMRI space 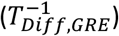 (Figure 3A). The main fibre direction was obtained by the 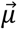 map from the NODDI analysis (Figure 3B).

**Figure 2:**
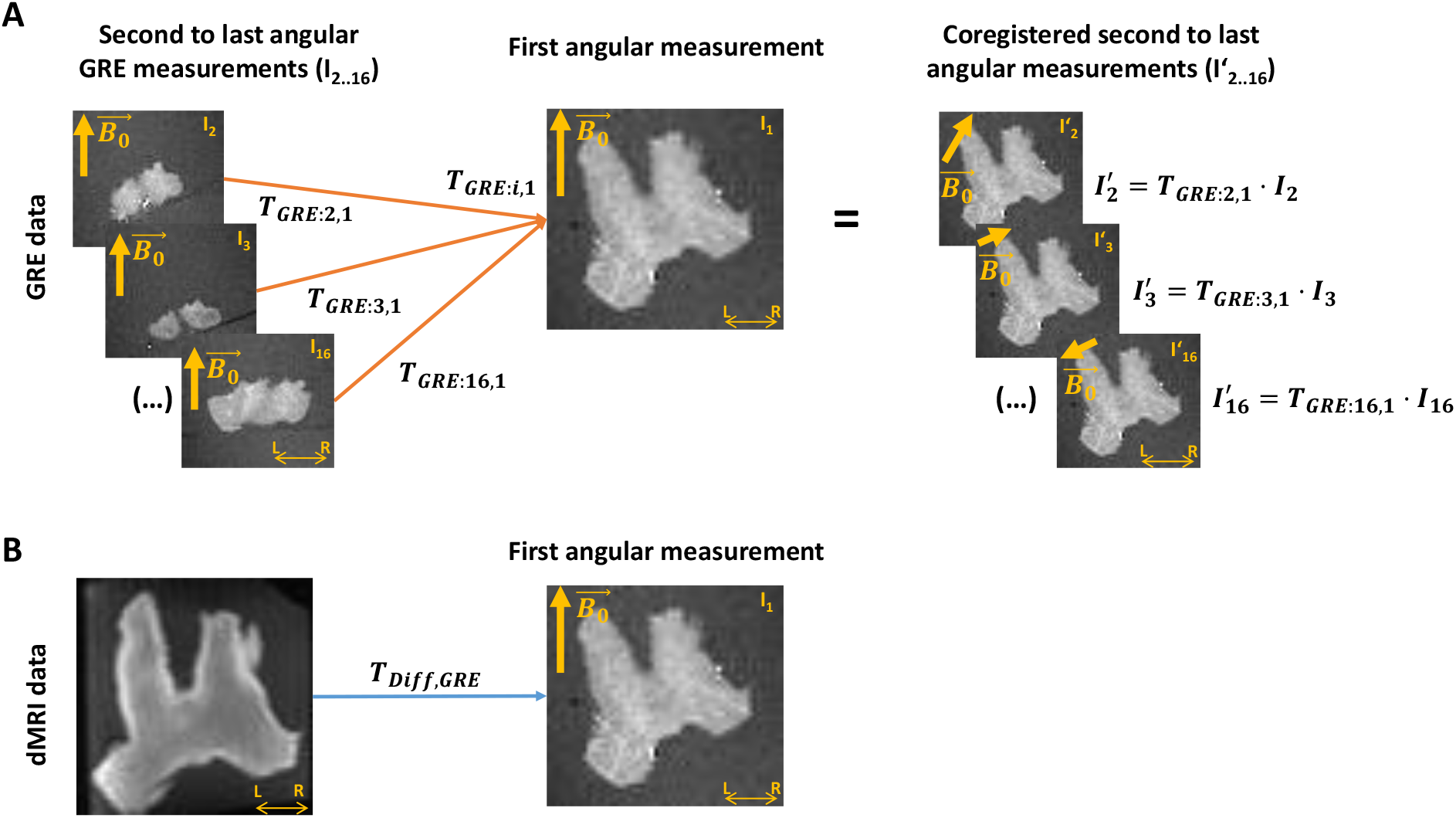
Coregistration of the ex vivo GRE and dMRI measurements. (A) A transformation matrix (T_GRE_) is obtained by coregistering all other multi-echo gradient-recall-echo (meGRE) datasets (I_2…16_) to the first measurement (I_1_, T_GRE: i,1_). This transformation matrices not only align, voxel-wise, the images of the meGRE datasets (I’_2…16_) to the first dataset, but also adjusts the directions of the main magnetic field 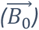 per angular measurement to preserves their relative orientation with respect to the first meGRE dataset. (B) A transformation matrix (T_Diff,GRE_) is obtained by coregistering the diffusion MRI (dMRI) image to the first angular GRE measurement. This transformation will allow the coregistration of the NODDI analysis results to the GRE data.

**Figure 3:**
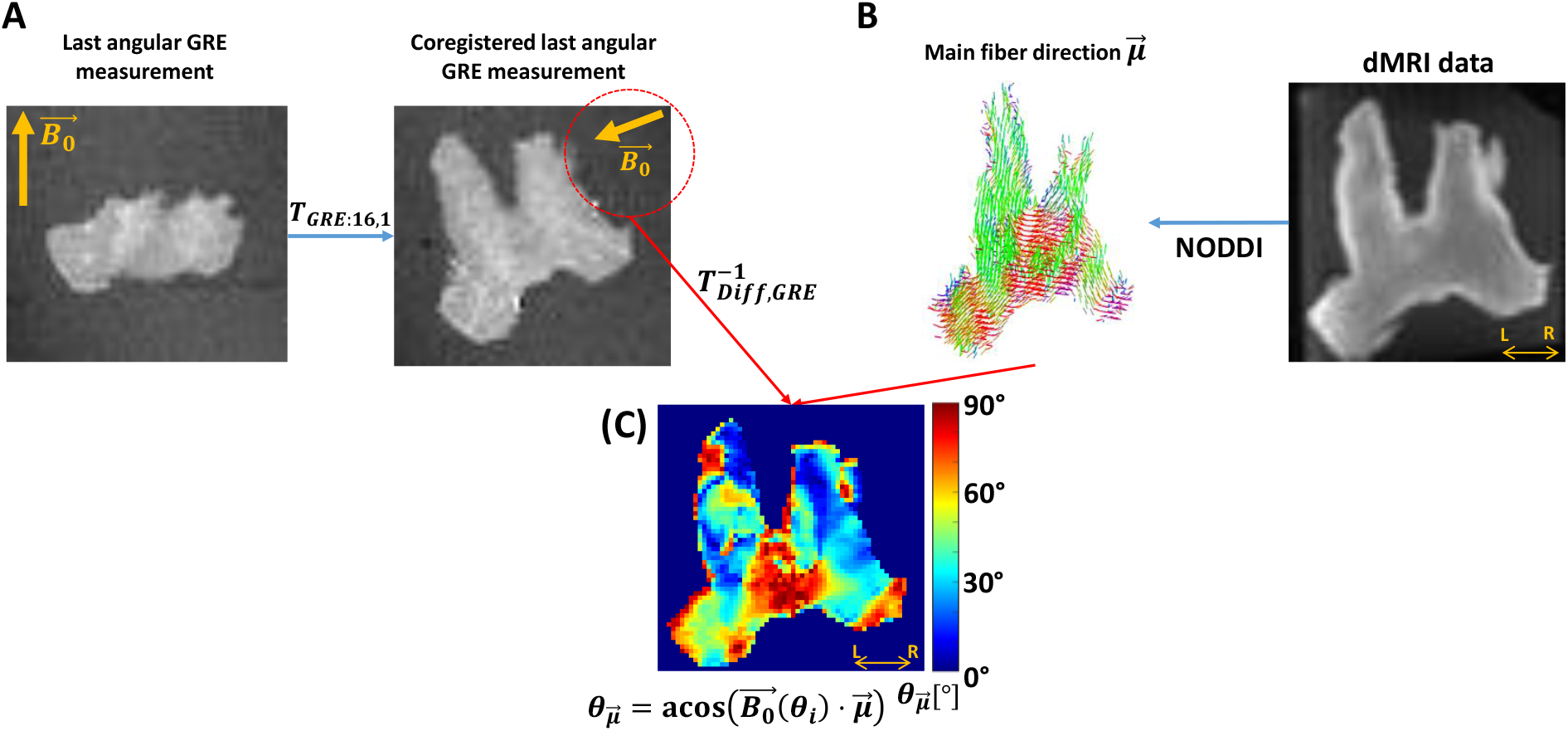
Estimation of the voxel-wise angular 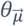 map. This estimation needed the B_0_ direction per angular GRE measurement 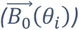 in diffusion space and the main fibre direction. (A) The 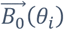 was estimated by applying to 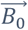, first, the transformation matrix between GRE volumes (T_GRE:i,1_) and later from GRE-to-diffusion (T^-1^_Diff,GRE_). (B) The main fibre direction 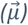 was acquired by analysing the dMRI data with the NODDI model. (C) Then, the voxel-wise 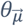 per angular measurement was computed by the arccosine of the scalar product between the projected 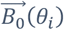 and the main diffusion direction 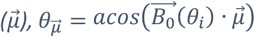. This sketch shows the steps for the last GRE angular measurement.

Note that 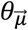 was computed in dMRI space instead of GRE space to avoid undersampling and interpolation because of transforming 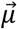 to GRE space. These sources of error do not occur by transforming 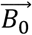 to dMRI space, i.e. computing 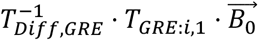, for each GRE angular measurement, since it is a global rather than a per-voxel measure. Finally, the 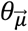 maps together with the ICVF and κ maps (not shown in Figure 3) were transformed using *T*_*Diff,GRE*_. Exemplary 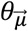 maps in GRE space are shown in Supplementary Material, Figure S1 (first row).

#### 3.1.5. Masking and pooling the *ex vivo* data

Before analysis, the *ex vivo* data required further pre-processing to remove outliers and to ensure a robust assessment of the effect of fibre dispersion and 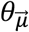 on R_2_*. For that, the *ex vivo* data were masked using the coregistered ICVF map and later pooled across the sixteen coregistered meGRE angular measurements.

In this process, all voxels in the OC with an ICVF > 0.8 were selected and pooled across all the meGRE angular measurements, hereafter referred to as cumulated data. The ICVF threshold was used because the extra-axonal space in the *ex vivo* specimen is reduced, e.g. (Stikov et al., 2011). The application of this threshold reduced the number of voxels in the OC by a 7.2% (∼ 600 over 8744 voxels). By pooling the data, the resulting cumulated data has the signal decays as a function of TE but also of 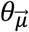, and fibre dispersion assessed by κ.

### 3.2. Simulated R _2_ * signal decay from the HCFM

R_2_* signal decay was simulated as ground truth (hereafter, *in silico* data) to assess the impact on M2 of variable fibre orientation, dispersion and myelination (i.e. g-ratio). For that, an averaged MR signal was calculated from an ensemble of 1500 hollow cylinders. The cylinders were evenly distributed on a sphere with defined spherical coordinates: an azimuthal angle ϕ rotating counter-clockwise from 0° to 359° starting at +X axis, and elevation angle θ rotating from 0° (+Z) to 180° (-Z). The signal contribution per hollow cylinder was modelled with the hollow cylinder fibre model (HCFM) for all the compartments, S_C_ (Equation A1).

In this work, two considerations were taken. First, the 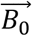 was fixed and oriented parallel to +Z (Figure 4A). Second, the approximated piece-wise function (Equation A8) of D_E_ (Equation A5) in the S_E_ signal (Equation A2b) was replaced by its analytical solution (Equation A9), because a discontinuity in this piece-wise function was observed in the so-called critical time (Wharton and Bowtell, 2013; Yablonskiy and Haacke, 1994). See section 9.2 for a detailed discussion.

**Figure 4:**
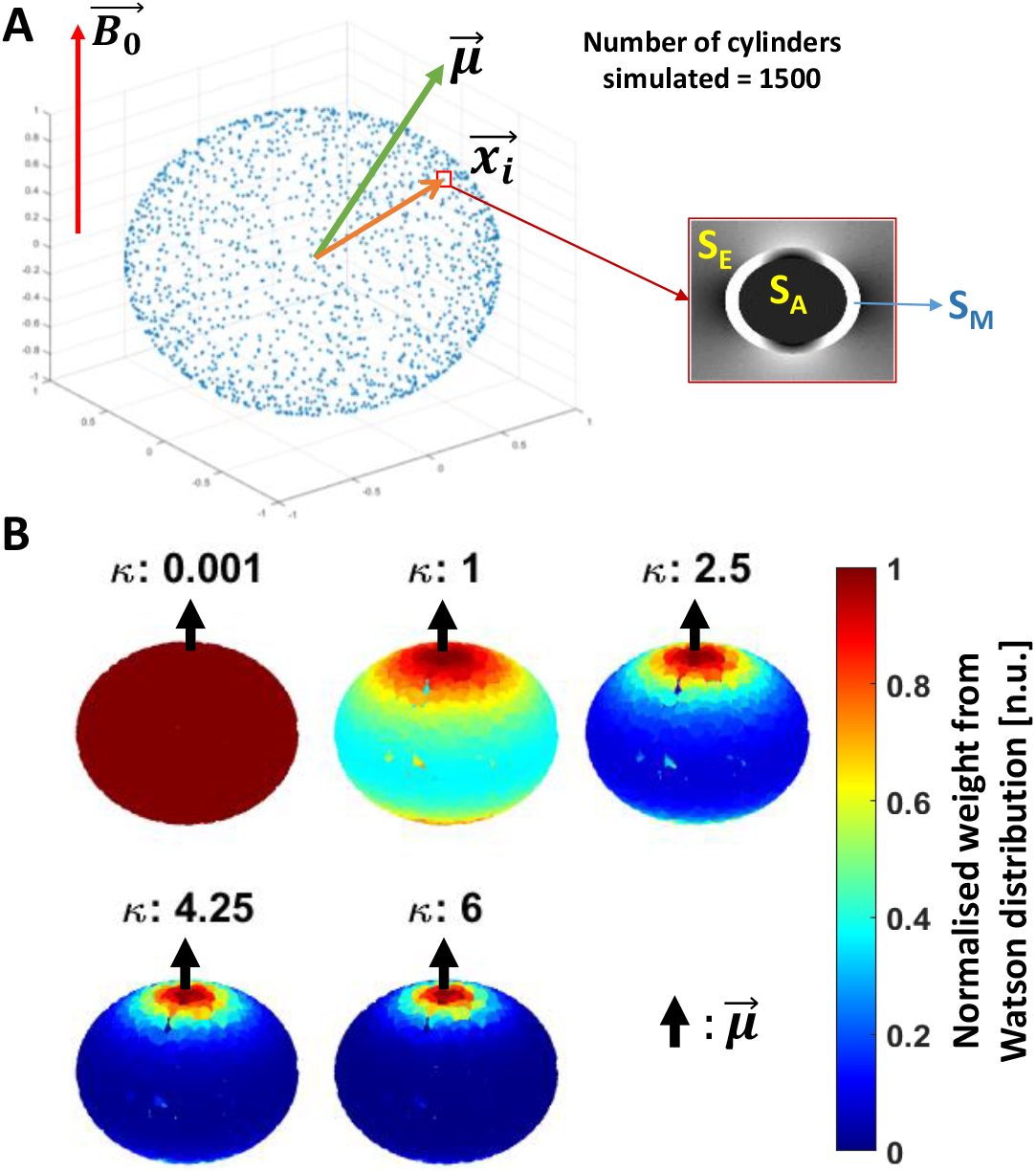
Schematics of the simulated in silico data: (A) Simulation: 1500 hollow cylinders, each of them defined by the vector 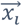, were distributed evenly on a sphere (see the blue dots). A mean orientation 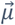 of the cylinders is defined, with the external magnetic field 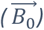 oriented parallel to the Z-axis. The signal contribution per cylinder was modelled using the Hollow Cylinder Fibre Model (HCFM) with the intra-axonal (S_A_), extra-axonal (S_E_) and myelin (S_M_) compartments (inset). (B) Addition of cylinder’s dispersion: the dispersion effect was added by weighting the signal coming from the cylinders by the parameter κ from the Watson distribution and 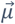 (Equation 9b). The parameter κ is limited from κ = 0 for isotropically dispersed to κ = infinity to fully parallel fibres. Here, 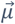 is parallel to 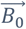.

To incorporate the effect of fibre dispersion in the *in silico* data, the ensemble-average signal was calculated by weighting S_c_ with the Watson distribution (W, (Sra and Karp, 2013) and Equation 6b). This weight from the Watson distribution was calculated using the position of each simulated cylinder, 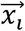, and a mean fibre orientation 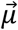, both defined with spherical coordinates (ϕ, θ) and 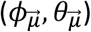, respectively. For simplification, 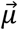 was restricted to an azimuthal angle of zero 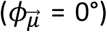. Then, the analytical expression of the ensemble-average signal, S_W_, is defined as follows:

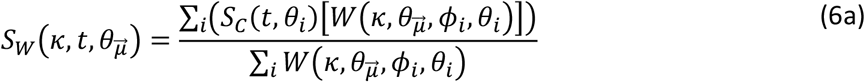

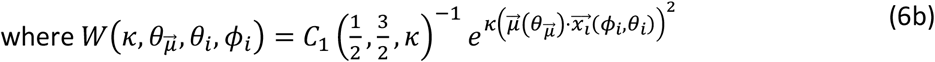

In Equation 9b, *C*_1_() is the confluent hypergeometric function, which is the normalisation factor of the Watson distribution, and the exponent holds the norm of the inner product between each individual i-th cylinder 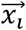 and 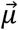. The level of dispersion was modulated by the parameter κ (Sra and Karp, 2013; Zhang et al., 2012) as shown in Figure 4B for a few cases. It is important to note that the notation 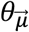 for the elevation angle of 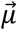 used here is equal to the one used to describe the fibre’s angular orientation in the *ex vivo* data (section 3.1.4). This is intentional since they stand for the same concept for both datasets. This simulation approach was used in previous conference publications (Fritz et al., 2020) and (Fritz et al., 2021).

With the ensemble averaged signal equation (Equation 6a), R_2_*-weighted signal decay can be created. In this work, the R_2_*-weighted signal decay was dependent on 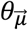, κ and g-ratio, as a function of time. The values used are reported in Appendix (Table A3). The remaining fixed parameters required by the ensemble-averaged equation were obtained from (Dula et al., 2010a; Wharton and Bowtell, 2013) and are listed in section 9.4 in Appendix (Table A1 and A2).

Finally, each simulated R_2_*-weighted signal decay was replicated 5000 times with an additive Gaussian complex noise (Gudbjartsson and Patz, 1995) to approximate the SNR of the experimental *ex vivo* data (see section 9.4). The experimental SNR was calculated by dividing the MR signal acquired at the first echo by the standard deviation of the background voxels of its corresponding image (Kellman and McVeigh, 2005), resulting in a mean SNR across the selected voxels of the OC of 112 (section 3.1.4).

### 3.3. Data analysis

#### 3.3.1. Data fitting and binning

The *ex vivo* data (section 3.1) and *in silico* data (each of 5000 replicas per simulated R_2_*-weighted signal decay, section 3.2) were analysed with the log-linear and log-quadratic models, M1 (Equation 2) and M2 (Equation 1), respectively. In both models, the *α*’s (*α*_0_ in arbitrary units, *α*_1_ in units of 1/s) from M1, and *β*’s (*β*_0_ in arbitrary units, *β*_1_ in units of 1/s and *β*_2_ in units of 1/s^2^) from M2, hereafter referred to as the *α*-parameters and *β*-parameters, were fitted as a function of TE. To fit the data, ordinary Least Square (OLS) optimization was used for both models in a custom-made Matlab code. Three fittings were performed at three different maximum TE values: TE_max_ = 54 ms (all 16 time points), TE_max_ = 36 ms (first 10 points) and TE_max_ = 18 ms (first 5 time points). Exemplary *α*_1_ and *β*_1_ maps obtained by fitting at TE_max_ = 54 ms on the *ex vivo* specimen are shown in Supplementary Figure S1 (middle and bottom row).

To compare the *α*- and *β*-parameters between datasets as a function of fibre dispersion (κ) and 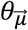, the fitted parameters were binned and averaged for the *ex vivo* cumulated data (section 3.1.5) and the *in silico* data. However, the *in silico* data required two extra averages on the fitted parameters: first, across the 5000 replicas and, second, across the κ values used for simulation. The average across κ was performed in such a way that it resembled the frequency distribution of κ observed in the *ex vivo* cumulated data (for more detail, see section 9.4).

In the binning process, both datasets were distributed first as a function of κ, and later as a function of 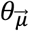. The first distribution was performed to ensure a similar degree of fibre dispersion as observed in Figure 4B and in the work of (Fritz et al., 2020). For that, three different fibre dispersion ranges were defined as a function of κ: κ < 1 for the highly dispersed fibres, 1 ≤ κ < 2.5 for the mildly dispersed fibres, and κ ≥ 2.5 for the negligibly dispersed fibres. Coincidentally, these fibre dispersion ranges depicted specific areas in the OC (Figure 5A).

**Figure 5:**
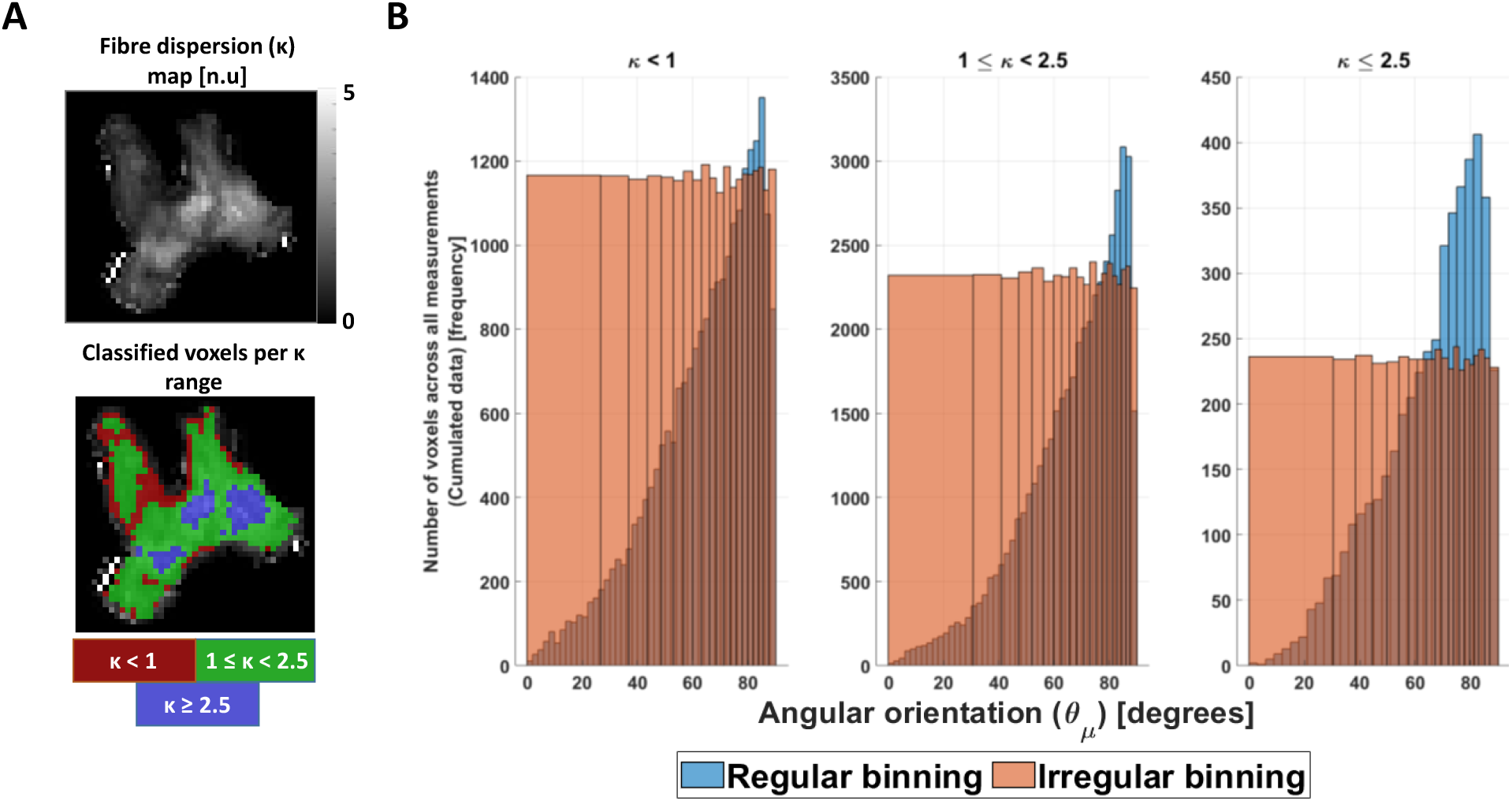
Preparation of the ex vivo data for analysis. (A) The cumulated ex vivo data was distributed first as a function of κ parameter, to ensure similar fibre dispersion. Heuristically it was divided in highly dispersed (κ < 1), mildly dispersed (1 ≤ κ < 2.5) and negligibly dispersed (κ ≥ 2.5) fibres. Coincidentally, this division enclosed specific areas in the OC (red, green and blue ROIs). (B) After division, the cumulated data were binned irregularly as a function of the estimated voxel-wise angular orientation 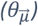 per κ range (orange bars), to avoid a possible effect size bias caused by its non-uniform distribution (blue bars). The first angular irregular bin or angular offset θ_0_ was obtained and showed to be κ range dependent (Table A4, section 9.4).

After separating the fitted parameters per fibre dispersion range for both datasets, the data was irregularly binned per 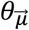 bin per defined κ range. This was performed to avoid any bias due to effect size, since a non-uniform distribution of voxels was found in the *ex vivo* cumulated data as a function of 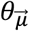 (Figure 5B, blue bars). To estimate the irregular 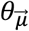 bin, a cumulated 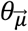 distribution of voxels was estimated and divided into 20 equally populated bins (Figure 5B, orange bars). The mean of the first angular irregular bin was defined as the angular offset *θ*_0_. The range of 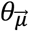 values contained in each irregular bin and the *θ*_0_ values are shown in Table A.4 in section 9.4.

After binning, the average and standard deviation (sd) was calculated per irregular 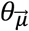 bin in the *ex vivo* cumulated data. For the *in silico* data, the average and sd were obtained by weighting the distribution of 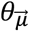 in each bin in a similar way to that seen in the irregular bins in the *ex vivo* cumulated data (for more detail, see section 9.4).

#### 3.3.2. Quantitative analysis

Four different analyses were performed for both datasets in order to study: (1) the effect of g-ratio and fibre dispersion, via κ, on the estimated angular-independent *β*_1_ using M2, (2) the microstructural interpretability of *β*_1_ via the deviation between fitted *β*_1_ and its predicted counterparts from M2 (*β*_1,nm_, Equation 3) and from the heuristic expression (*β*_1,m_, Equation 4), (3) the possibility of calculating the MWF and g-ratio from the fitted *β*_1_ using the heuristic expression *β*_1,m_ (Equation 4), and (4) the effect of TE on the performance of M2 in estimating R_2,iso_* from *β*_1_. The first two analyses were aimed to test whether *β*_1_ can be used as a proxy of R_2,iso_*.

For the first analysis, the capability of M2 to estimate an orientation-independent effective transverse relaxation rate, R_2,iso_*, via the *β*_1_ parameter was assessed. Since R_2,iso_* by definition is the angular independent part of R_2_ * and according to the HCFM should be given by *β*_1_ parameter at 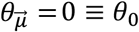, we assessed the residual 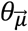 dependence of the *β*_1_ parameter with respect to *θ*_0_ and compared it with its counterpart for *α*_1_, i.e. the proxy for the 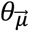 dependent R_2_*.

For this, we first calculated the 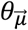 dependence of each parameter with respect to *θ*_0_ using the normalized-root-mean-squared deviation (nRMSD):

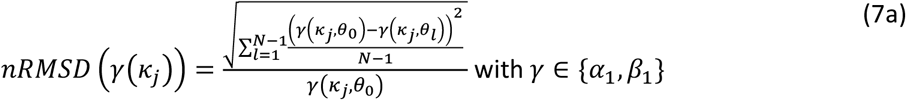

where *θ*_0_ varied slightly for each κ range (sub-index j) but was close to zero (see Table A.4 in section 9.4).

To compare the nRMSD of each parameter, we calculated the difference between them, ΔnRMSD, as:

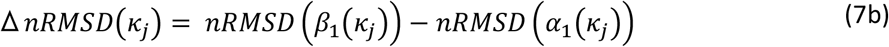

in percentage-points (%-points). If the ΔnRMSD is positive or higher than 0 %-points, this implies that the 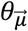 dependency of *β*_1_ is similar or higher, in magnitude, to *α*_1_. The latter says therefore that M2 failed in estimating an angular-independent parameter, or disentangling the 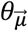 dependency from R_2_*. A negative ΔnRMSD in turn implies that the 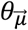 independence of *β*_1_(*κ*_*j*_) has been reduced. A perfect orientation independence is achieved if *nRMSD* (*β*_1_(*κ*_*j*_)) = 0 and, consequently, Δ*nRMSD*(*κ*_*j*_) = −*nRMSD* (*α*_1_(*κ*_*j*_)).

For the second analysis, the microstructural interpretation of *β*_1_ was quantitatively assessed by comparing the relative difference (ε) between estimated *β*_1_ at the angular orientation 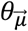 for the fitted *in silico* data 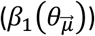 and the predicted *β*_1_ equivalence (*β*_1,*p*_) using M2 (Equation 3) or the heuristic expression (Equation 4):

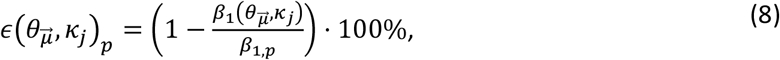

Where *p* ∈ {*nm, m*} and *β*_1,*nm*_ and *β*_1,*m*_ as defined in Equation 3 and 4. Additionally, the mean 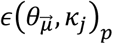 across angles was calculated as 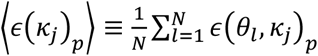.

The above analyses were performed per g-ratio across κ range using the same values as for *in silico* data (Tables A1, A2 and A3, section 9.4). For the third analysis, *β*_1,m_ (Equation 4) was rearranged to estimate MWF from the fitted *β*_1_ in *ex vivo* data. For that, the R_2_ values of the non-myelinated (R_2N_) and myelinated (R_2M_) compartments are reported in Table A1. After estimating the MWF, the g-ratio values were also estimated by rearranging Equation 5. For that, the fibre volume fraction, FVF, and proton density values, ρ_N_ and ρ_M_, required for this calculation are reported in Table A1.

For the last analysis, the effect of TE on the capability of M2 to estimate R_2,iso_* via *β*_1_ was assessed by comparing the *ex vivo* dataset and the *in silico* data with similar g-ratio, obtained from the previous analysis. For that, *α*_1_ and *β*_1_ from M1 and M2 were compared once again as in the first analysis. However, now the models were fitted to meGRE datasets with different longest echo times (TE_max_): 54 ms, 36 ms and 18 ms. Again, the ΔnRMSD was calculated to assess the residual 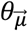 dependence of *β*_1_ in comparison to the 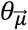 dependence of *α*_1._

In the following sections, the dependency of the parameters under study, i.e. 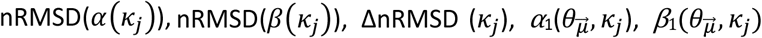 (Equations 7a-b), 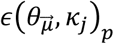 (Equation 8) and ⟨*ϵ*(*κ*_*j*_)_*p*_⟩, to 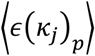, to 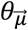 and *κ* were simplified for readability purposes. Therefore, the aforementioned parameters will be hereafter nRMSD(*α*_1_), nRMSD(*β*_1_), ΔnRMSD, *α*_1_, *β*_1_, *ϵ*_*p*_ and ⟨*ϵ*_*p*_⟩, respectively.

## 4. Results

### 4.1. First analysis: Capability of M2 to obtain the angular-independent *β*_1_ parameter for varying g-ratio and fibre dispersion values

Figure 6 and 7 show the capability of M2 to estimate R_2,iso_* via *β*_1_ for variable g-ratio and fibre dispersion. To visualise this, the 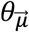 dependency of *α*_1_ from M1 was compared to the residual 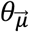 dependency *β*_1_ from M2 (Figure 6A and 6B). Both 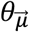 dependencies were quantified in Figure 7A and 7B using their respective nRMSD (Equation 7a) and the ΔnRMSD in Figure 7C (Equation 7b). The results are from the analysis performed on the *in silico* and *ex vivo* data.

**Figure 6:**
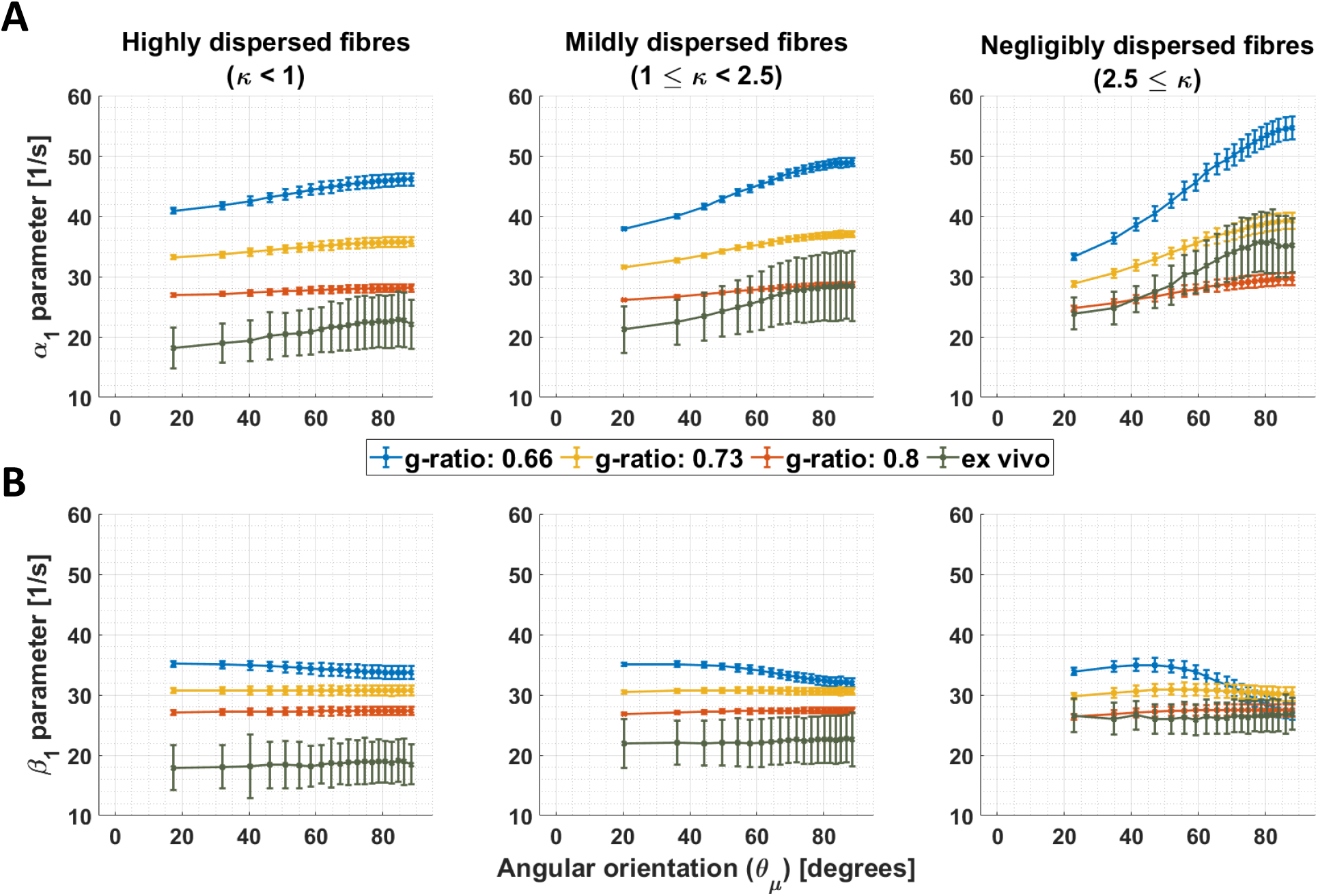
Orientation dependence of linear model parameters (α_1_ and β_1_) for varying g-ratio and fibre dispersion values. (A-B) Depicted is the α_1_ parameter of M1 (proxy for R_2_*) and β_1_ parameter of M2 (proxy for the isotropic part of R_2_*) as a function of the angle between the main magnetic field and the fibre orientation 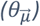 for different fibre dispersion and g-ratio values. The different columns depict different dispersion regimes: highly dispersed (κ < 1, first column), mildly dispersed (1 ≤ κ < 2.5, second column) and negligibly dispersed (κ ≥ 2.5, third column) fibres. The distinct colours distinguish between in silico data with variable g-ratios (0.66 in blue curve, 0.73 in yellow curve, and 0.8 in red curve) and ex vivo data (olive curve). Note that the smallest angle (θ_0_) varied across dispersion regimes: 17.3° (κ < 1), 20.4° (1 ≤ κ < 2.5) and 22.9° (2.5 ≤ κ). This was caused by the irregular binning (see section 3.1.4).

**Figure 7:**
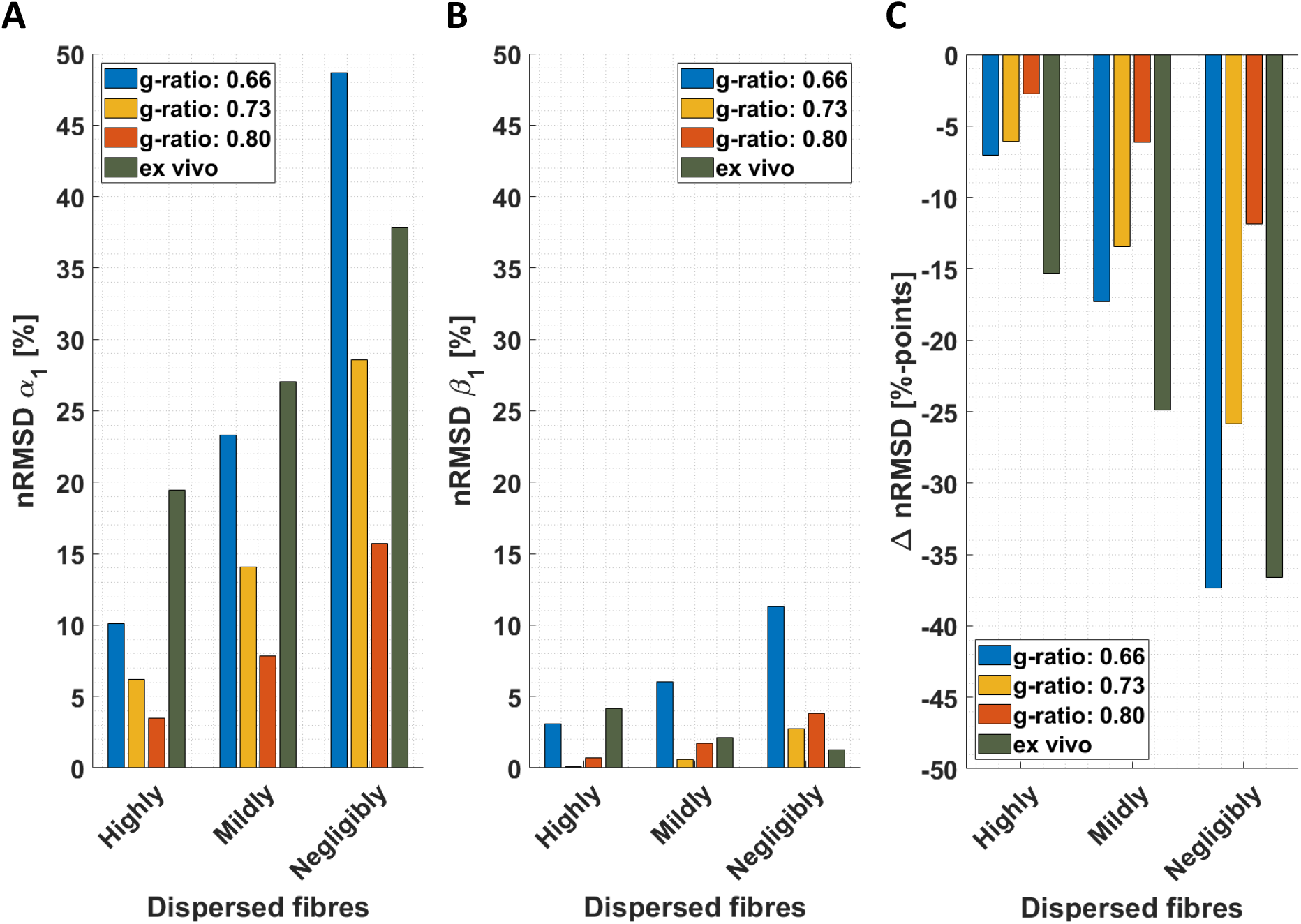
Quantifying orientation dependence of linear model parameters (α_1_ and β_1_) for varying g-ratio and fibre dispersion values. (A-B) Depicted is the normalised root-mean-squared deviation (nRMSD, Equation 7a in %) of the α_1_ parameter of M1 (proxy for R_2_*) and β_1_ parameter of M2 (proxy for the isotropic part of R_2_*) for different fibre dispersion and g-ratio values. (C) Depicted is ΔnRMSD (Equation 7b in % points) comparing the residual 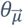 dependency of β_1_ with the 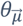 dependency of α_1_ (negative values mean M2 reduced the 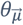 dependency of R_2_*). The four coloured bars (i.e. [blue, red, yellow, olive]) per dispersion ranges (highly dispersed, κ < 1; mildly dispersed, 1 ≤ κ < 2.5; and negligible dispersed, κ ≥ 2.5 fibres) distinguish between in silico data with variable g-ratios (0.66 in blue bar, 0.73 in yellow bar, and 0.8 in red bar) and ex vivo data (olive bar).

The capability of M2 to reduce the 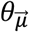 dependency of *β*_1_ varied with g-ratio and fibre dispersion, the 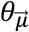 dependency of *α*_1_ was also strongly influenced by g-ratio and fibre dispersion: smaller g-ratio values and reduced fibre dispersion increased the 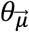 dependency of *α*_1_ and (the residual 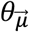 dependency) of *β*_1_ (Figure 6A and 6B, respectively).

The fibre dispersion affected the performance of M2 differently between *in silico* and *ex vivo* datasets. For the *ex vivo* data, the nRMSD(*β*_1_) was the lowest for the negligibly dispersed fibres (nRMSD(*β*_1_): 1.3% at κ ≥ 2.5) but less so for the highly dispersed fibres (nRMSD(*β*_1_) : 4.1% at κ < 1). For the *in silico* data, the nRMSD(*β*_1_) was the lowest for the highly dispersed fibres and for a g-ratio of 0.73 (nRMSD(*β*_1_): 0.1% to 2.7% with decreasing fibre dispersion). The nRMSD(*β*_1_) was higher, but still below 12%, for g-ratios of 0.66 and 0.8. The 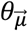 dependency of *α*_1_ on fibre dispersion was the same between *in silico* and *ex vivo* datasets: the lower the dispersion the higher the nRMSD(*α*_1_). The 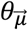 dependency of *α*_1_ increased the lower the g-ratio was. When comparing the residual 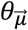 dependency of *β*_1_ with the 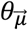 dependency of *α*_1_, the improvement is large for negligible dispersion (from ΔnRMSD = -11.9%-points to ΔnRMSD = -37.4%-points) for both datasets.

### 4.2. Second analysis: Assessment of the microstructural interpretability of *β*_1_

Figure 8A and 8B report the angular-orientation 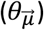 dependent relative differences (*ϵ*_*nm*_ and *ϵ*_*m*_, Equation 8) between the fitted *β*_1_ from the *in silico* data and its predicted counterparts using M2 (Equation 3) and the heuristic expression (Equation 4). Figure 8C shows the mean and standard deviation of *ϵ*_*nm*_ and *ϵ*_*m*_ across angles.

**Figure 8:**
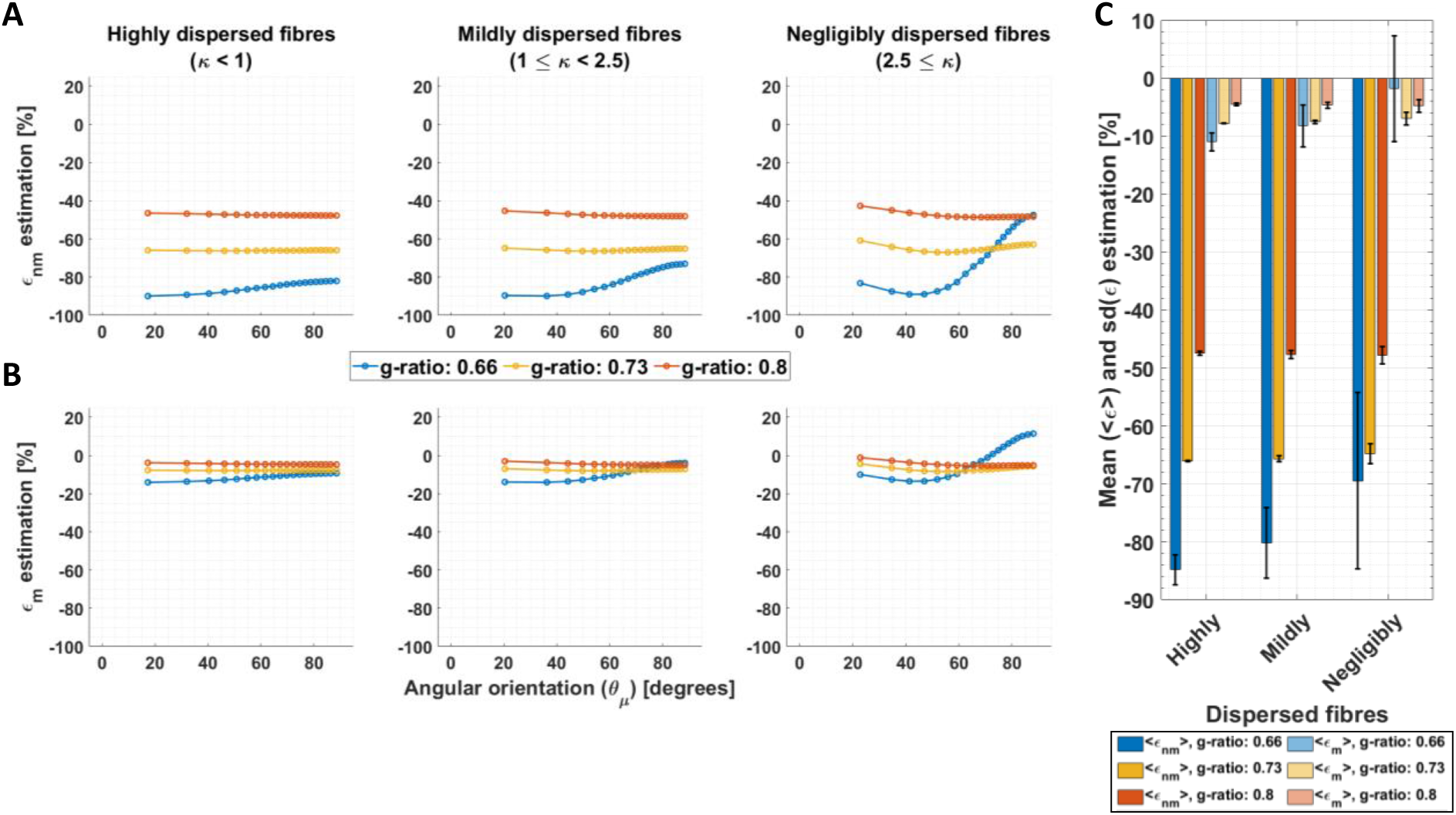
Assessment of the microstructural interpretability of β_1_ by the deviation between fitted and biophysically predicted β_1_. The relative difference (ε, Equation 8) was calculated between the fitted β_1_ to the in silico data and two biophysically-modelled expressions for β_1_ based on the HCFM. The two expressions for β_1_ values were calculated from the original expression for M2, β_1,nm_ (Equation 3, resulting in ε_nm_) and the heuristic expression, β_1,m_ (Equation 4, resulting in ε_m_). This was calculated per g-ratio and fibre dispersion. (C) The corresponding mean, <ε>, and standard deviation, sd(ε), of the relative differences across the angular orientations 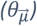 were estimated.

*ϵ*_*nm*_ was large, between -100% and -40%, and varied strongly with g-ratio and fibre dispersion. Even more, *ϵ*_*nm*_ showed an 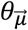 dependence where the largest deviation was observed for the smallest g-ratio (0.66) and the lowest fibre dispersion (Figure 7A). By contrast, *ϵ*_*m*_ was smaller, between -20% and 20%, and showed a smaller 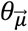 dependence, which was largest for the smallest g-ratio and lowest fibre dispersion. On average, we found that negligibly dispersed fibres showed the smallest *ϵ*_*nm*_ and *ϵ*_*m*_.

The mean across angles for *ϵ*_*nm*_, ⟨*ϵ*_*nm*_⟩, was smaller than 85% whereas the mean across angles for *ϵ*_*m*_, ⟨*ϵ*_*m*_⟩,was smaller than 12% (Figure 8C). On average across all g-ratios and fibre dispersion arrangements, ⟨*ϵ*_*nm*_⟩ was approximately 8 to 9 times larger than ⟨*ϵ*_*m*_⟩. Both relative mean differences decreased with increasing g-ratio and decreasing fibre dispersion for almost all ⟨*ϵ*_*nm*_⟩ and ⟨*ϵ*_*m*_⟩. ⟨*ϵ*_*m*_⟩ for the negligibly dispersed fibres at g-ratio 0.66 was close to 2% but accompanied by a large standard deviation across 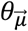, indicating strong 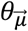-dependency of the corresponding fitted *β*_1_ parameters. For both *ϵ*_*nm*_ and *ϵ*_*m*_, the variability (Figure 8C) across different 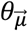 values, *sd*(*ϵ*_*nm*_) and *sd*(*ϵ*_*m*_) respectively, was highest when the fibre dispersion and g-ratio were lowest.

### 4.3. Third analysis: Myelin water fraction (MWF) and g-ratio estimation from *ex vivo* data using the heuristic expression of R_2,iso_* via *β*_1,m_

Figure 9 reports the MWF estimated from the *ex vivo* data by inverting the heuristic expression for *β*_1,m_ (Equation 4). Figure 9A shows the estimated MWF as a function of 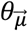 while Figure 9B shows the median and standard deviation (sd) of the estimated MWF across 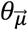.

**Figure 9:**
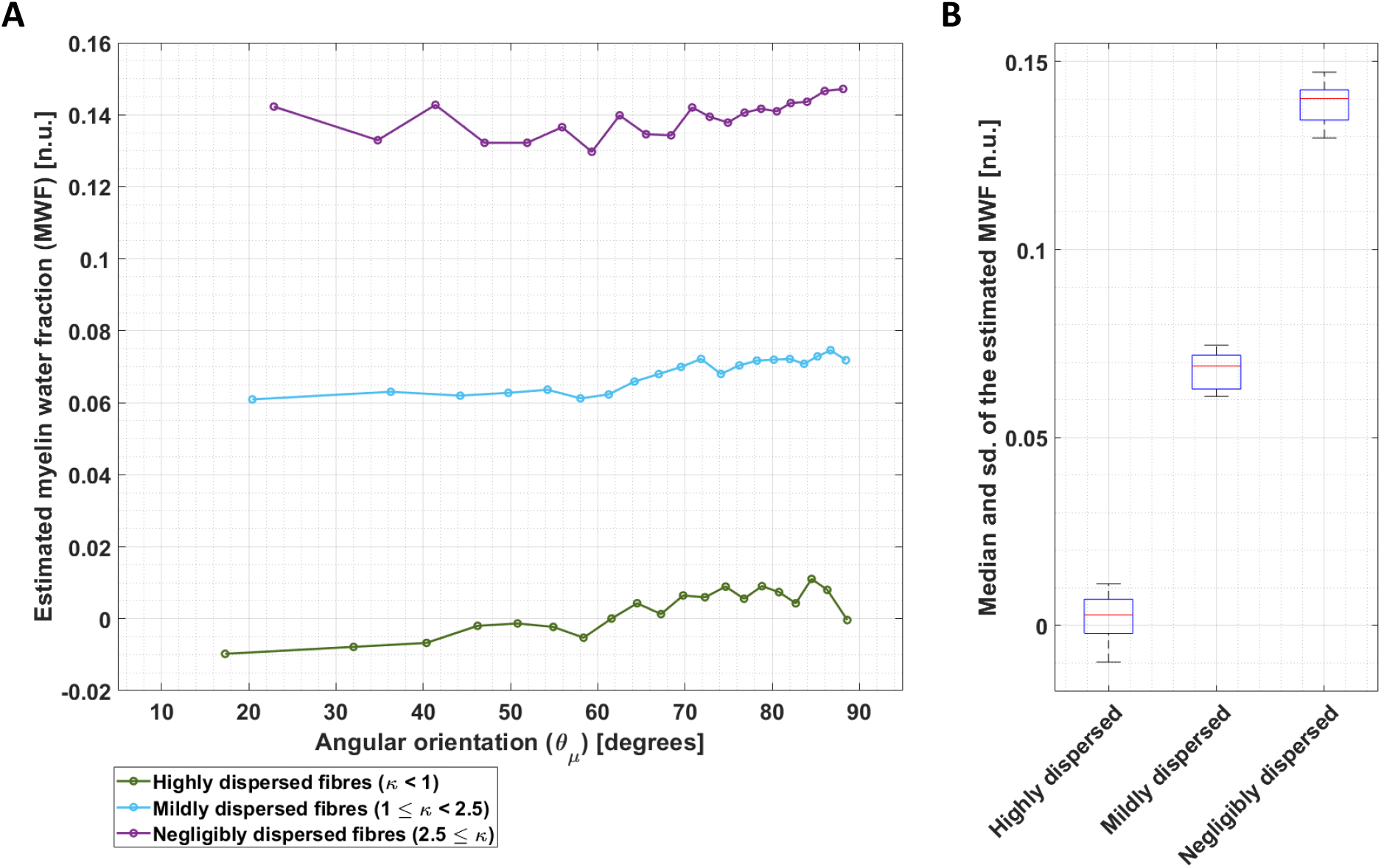
Dependence of the MWF estimation on angular orientation for three different fibre dispersion ranges in ex vivo data. (A) The MWF was estimated by using the heuristic analytical expression of β_1_ (β_1,m_, Equation 4) and the fitted β_1_ for the ex vivo data using the compartmental R_2_ values in Table A2. This calculation was performed per angle 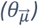 and for the three different fibre dispersion ranges: highly dispersed (green olive), mildly dispersed (cyan) and negligibly dispersed (magenta). (B) The corresponding median and standard deviation (sd) were estimated across 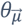 per fibre dispersion range.

The estimated MWF was larger with decreasing fibre dispersion (Figure 9A). Moreover, there was a trend towards larger estimated MWF for larger 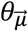. On average across 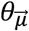 (Figure 9B), the estimated median *ex vivo* MWF decreased by 98% from highly dispersed fibres (MWF: 0.0028) and by 50.8% from mildly dispersed fibres (MWF: 0.069) in comparison to negligibly dispersed fibres (MWF: 0.14). The standard deviation across MWF was similar for different fibre dispersions, ranging from 0.0068 to 0.0104.

With the estimated median MWF, a median g-ratio can be estimated using Equation 5. The estimated median g-ratios were 0.996, 0.895 and 0.785 for highly, mildly and negligibly dispersed fibres, respectively.

### 4.4. Fourth analysis: the effect of echo time on the performance of M2

Figures 10A and 10B show the 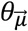 dependency of *α*_1_ and *β*_1_ as a function of TE_max_ for the *ex vivo* data and the *in silico* data, for the negligibly dispersed fibres (i.e., κ ≥ 2.5). Figures 11A and 11B show the corresponding nRMSD (Equation 7a) for both parameters at different TE_max_, while Figure 11C shows the difference between both nRMSD (ΔnRMSD, Equation 7b). Note that only negligibly dispersed fibres and the *in silico* data at g-ratio of 0.8 are studied here, because it is known from the results in Figure 8 that those possess the smallest relative difference and sd to the model predictions.

**Figure 10:**
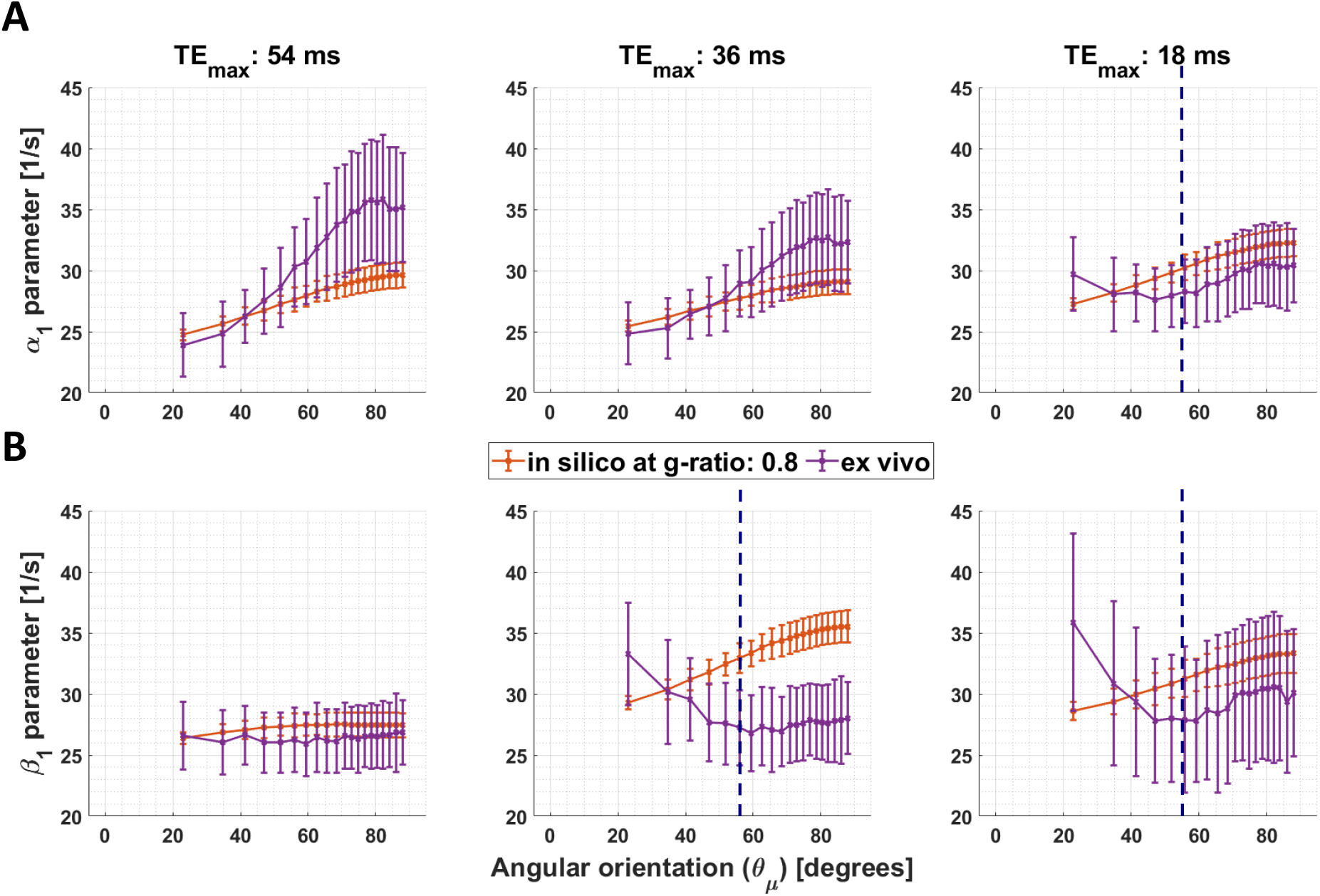
Effect of the maximal echo time on the 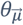 dependency of α_1_ and β_1_. (A and B) Angular orientation 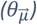 dependence of α_1_ in M1 and β_1_ in M2 for varying maximum TE (TE_max_: 54 ms, 36 ms and 18 ms). Two datasets are compared: ex vivo (magenta curve) and in silico (red curve) data at g-ratio of 0.8 which is closest to the estimated g-ratio of the ex vivo data. Moreover, only datasets of the negligibly dispersed fibres (κ ≥ 2.5) are presented. The blue vertical lines in some of the subplots indicates the magic angle 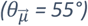.

**Figure 11:**
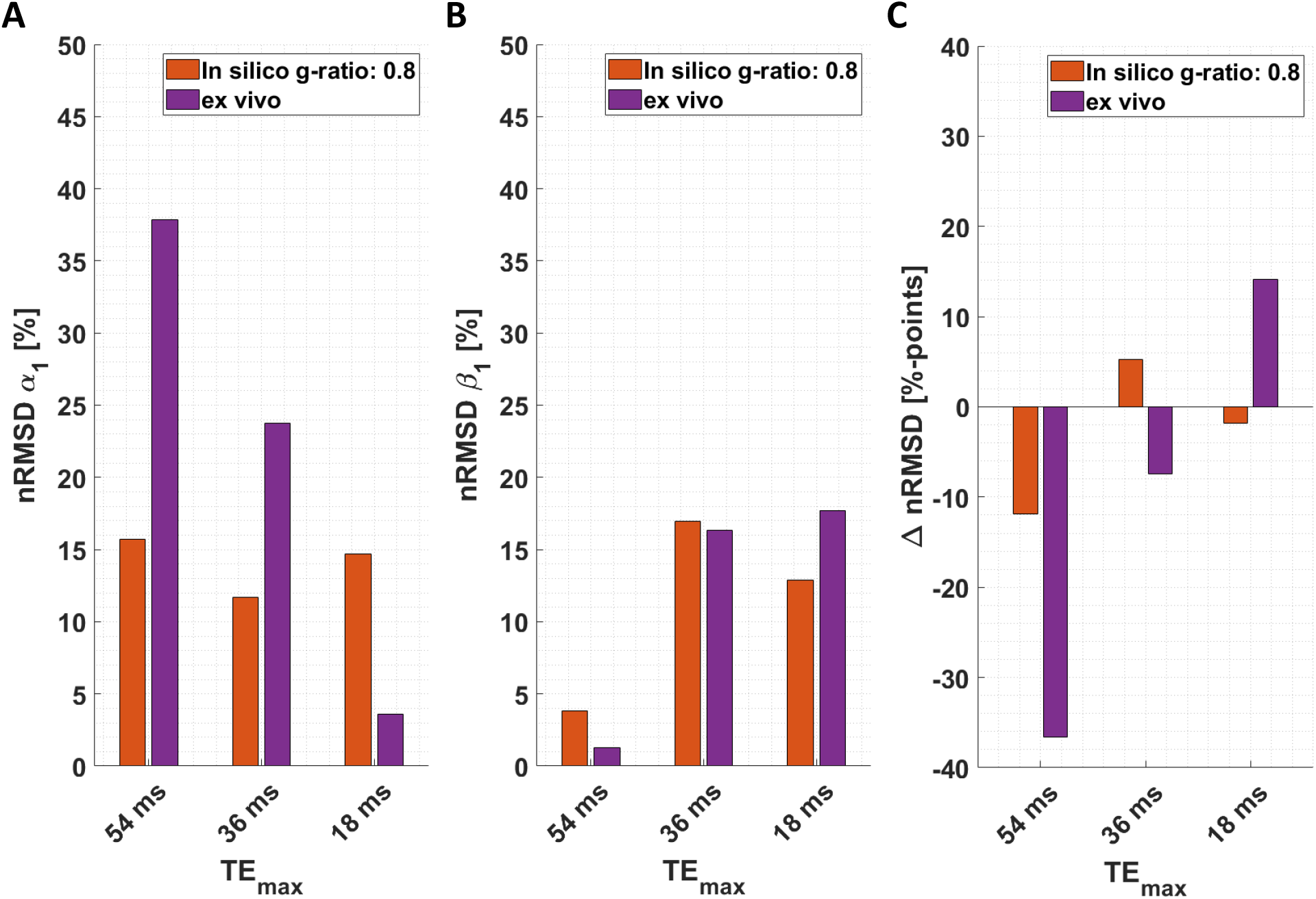
Quantifying the effect of the 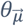 dependency of α_1_ and β_1_ for three different maximal echo times (TE_max_). (A-B) Depicted is the normalised root-mean-squared deviation (nRMSD, Equation 7a in %) of the α_1_ parameter of M1 (proxy for R_2_*) and β_1_ parameter of M2 (proxy for the isotropic part of R_2_*) for different TE_max_ values (54 ms, 36 ms and 18 ms), respectively. The distinct colours distinguish between in silico data at g-ratio of 0.8 (red bar) and ex vivo data (magenta bar), both for negligible dispersed fibres (κ ≥ 2.5). (C) Depicted is ΔnRMSD (Equation 7b in %-points) comparing the residual 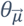 dependency of β_1_ with the 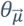 dependency of α_1_ (negative values mean M2 reduced the 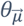 dependency of R_2_*),

At the largest TE_max_ *ex vivo* and *in silico* data showed the same trend (Figure 10, first column). M2 could greatly reduce the 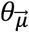 dependency of *β*_1_ when compared to the 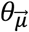 dependency of *α*_1_ (Figure 11A-B): nRMSD(*α*_1_) of 15.7% (*in silico*) and 37.9% (*ex vivo*) was reduced to 3.8% (*in silico*) and 1.3% (*ex vivo*). At smaller TE_max_ (36 ms and smaller), M2 was less effective (Figures 10, second and third column). Even an increased 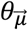 dependency was observed for *β*_1_ when compared to *α*_1_ (Figure 11C): ΔnRMSD = 5.3%-points at 36 ms (*in silico*) and ΔnRMSD = 14.1%-points at 18 ms (*ex vivo*). Moreover for the smallest TE_max_ (18 ms), an atypical 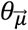 dependence of *β*_1_ (and *α*_1_) was found in the *ex vivo* data: *β*_1_ (and *α*_1_) decreased with increasing 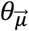 up to approximately 55° (magic angle, dashed blue lines in Figure 10A and 10B) and then slightly increased again. The 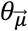 dependency up to the magic angle was not observed in the *in silico* data at any investigated TE_max_. Moreover, the 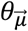 dependency of *α*_1_ in the *ex vivo* data decreased with decreasing TE_max_ also this trend was not observable in the *in silico* data (Figure 10A).

## 5. Discussion

This work quantitatively explored the efficiency of the log-quadratic model (M2) in deciphering the orientation-independent part of R_2_* (R_2,iso_*) via its linear parameter, *β*_1_, from a single-orientation multi-echo GRE (meGRE) while varying microstructural fibre properties, i.e. fibre dispersion and g-ratios. Our findings demonstrated that M2 was effective in estimating R_2,iso_* via *β*_1_ when using meGRE with long maximum echo time (TE_max_ ≈ 54 ms) for all investigated microscopic arrangements in both simulations and *ex vivo* measurements. Moreover, we confirmed our hypothesis that the fitted *β*_1_ of the meGRE signal from tissue with different g-ratios cannot be predicted using the biophysical relation in M2, which is derived from the hollow cylinder fibre model (HCFM) and neglects the contribution of the myelin water compartment. We proposed a heuristic expression that predicts *β*_1_ to be a weighted sum of the relaxation rates of the myelin and non-myelin water pools and showed that this expression can better describe the data. Using the heuristic expression we estimated the MWF and the g-ratio from the fitted *β*_1_ of the *ex vivo* data achieving plausible results. Lastly, we found that M2 was not capable of estimating R_2,iso_* correctly when using shorter maximum echo times (TE_max_ < 36 ms) that are typical of whole-brain *in vivo* protocols. We made another unexpected observation at the shortest investigated TE_max_ (18 ms): Here, the orientation-dependency of the classical R_2_* showed the highest deviation between *ex vivo* and *in silico* data for angles below the magic angle (55°) indicating that at short echo times the mechanism for this orientation-dependency of R_2_* is not captured by the HCFM-based simulation used here.

### Capability of M2 to estimate the angular independent *β*_1_ for varying g-ratio and fibre dispersion values

M2 has the potential to estimate R_2,iso_* from a single-orientation meGRE via *β*_1_. By assessing the residual 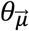 dependency of *β*_1_ we found that M2 was effective although its capability varied for different g-ratios and fibre dispersions (Figure 6). Nevertheless, the residual 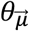 dependency of *β*_1_ was always less than 12% even if the 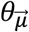 dependency of the original R_2_* (via the *α*_1_ parameter of M1) was up to 50% (Figure 7A). The residual 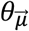 dependence of *β*_1_ was smallest at a g-ratio of 0.73 (Figure 7B). The highest performance of M2 was found for negligibly dispersed fibres at the lowest g-ratio (0.66, Figure 7C), where an original 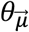 dependence of R_2_* (i.e. via the *α*_1_ parameter of M1) of almost 50% was reduced to less than 12% in the 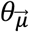 dependency of *β*_1_. Note that the g-ratio value at which the performance of M2 is maximal might also depend on the compartmental R_2_ values. In the simulations, we used the R_2_ values from (Dula et al., 2010b) (Table A2). It is possible that using different R_2_ values would result in a different g-ratio values for which the model’s performance was maximal.

### Assessment of the microstructural interpretability of *β*_1_

M2 is derived from the biophysical HCFM (Wharton and Bowtell, 2013, 2012) and thus the fitted *β*_1_ parameter can be related to microscopic tissue parameters (Equation 16b, section 9.3). We found an error of up to 85% (Figure 8C) between the *β*_1_ obtained by fitting M2 to the *in silico* data and the predicted *β*_1_ parameter using the biophysical relations in M2 (Equation 3). This confirms our hypothesis that neglecting the myelin contribution in the derivation of M2 results in an invalid biophysical expression for *β*_1_. We showed that the proposed heuristic expression for *β*_1_ is better suited for biophysical interpretation than the original version from M2, resulting in a relative error that was less than 12% (Figure 8C). The newly found heuristic expression for *β*_1_ implies that in the slow-exchange regime, the linear time-dependent component of the logarithm of the meGRE signal can be described as a sum of the relaxation rates of the myelin and non-myelin water pools weighted by their signal fractions (Equation 4). This can be understood when considering that the *β*_1_ parameter in M2 captures the contribution of the linear component of the logarithmic–signal decay, which is the equivalent to an effective mono-exponential decay. The effective relaxation rate of a mono-exponential decay, however, can be expressed as the sum of compartmental relaxation rates weighted by their corresponding signal fractions as is well-known from the fast exchange regime. In other words, we showed here that when approximating the logarithm of the signal of a two-pool model in the slow-exchange regime by a second order polynomial in time, the first-order term in time captures the fast-exchange regime behaviour of the signal decay whereas the second-order term in time accounts for deviations from the linear (i.e. mono-exponential) signal decay.

Note that the proposed heuristic correction does not account for the effect of fibre dispersion which might explain why the accuracy of the prediction was reduced with increasing fibre dispersion. While the influence of fibre dispersion has been successfully incorporated into M2 in another study (Fritz et al., 2020), it remains an open task for future studies to do this as well for the heuristic expression of *β*_1_.

### Myelin water fraction and g-ratio estimations from *ex vivo* data using the heuristic expression of *β*_1_

Assuming an effective performance of M2 in estimating the angular-independent *β*_1_ and using the heuristic expression for its biophysical interpretation, MWF and g-ratio can be estimated from the fitted *β*_1_ of the *ex vivo* data (Equation 5). For the negligibly dispersed fibres we found a median (across orientation) MWF of 0.14 (Figure 9B), which is congruent with the mean value reported in white matter of 0.10 (Uddin et al., 2019). By using the FVF and proton density values from the *in silico* data (Table A1), we found a median g-ratio of 0.79. The estimated g-ratio value is higher than typical MRI-based g-ratio values reported for the *in vivo* brain, which ranges between 0.65 and 0.70 (Berman et al., 2018; Emmenegger et al., 2021; Stikov et al., 2015) but is close to the value used in the work by (Wharton and Bowtell, 2013). The reason for the dissimilarity between predicted g-ratio and its counterpart from literature might be related to the additional assumptions that were made to estimate the g-ratio: while the estimation of MWF only requires knowledge of the compartmental R_2_ values, the g-ratio estimation requires additional knowledge of fibre volume fraction (FVF) and proton density values (Equation 6).

In this study, we used the same parameters as reported in (Wharton and Bowtell, 2013) for the FVF, and the proton densities, whereas the compartmental R_2_ values were based on (Dula et al., 2010b). Particularly, the value employed for the FVF (0.5, Table A1, section 9.4) is considerably lower than the values reported in literature, e.g. 0.75 in Stikov et al., 2011, which might explain the similarity between our reported g-ratio value and the one from (Wharton and Bowtell, 2013). Furthermore, we found that increasing the amount of fibre dispersion leads to a substantial underestimation of the MWF and an overestimation of the g-ratio. This might be of practical importance when using this method for MWF or g-ratio estimation.

### The effect of echo time on the performance of M2

We found that M2 is less effective in estimating R_2,iso_* when the maximum TE is reduced to values more typically used for *in vivo* studies (i.e., maximal TE of 18 ms) but also at intermittent echo time ranges (36 ms, Figure 11). This observation is at first glance in contradiction with the validity range of M2 because the mathematical approximations, especially the approximation of the dephasing component of the extra-cellular compartment D_E_ (section 9.2, Figure A1), are valid only in the regime TE ≤ 36 ms (see section 9.3). This apparent contradiction can be resolved by acknowledging the contribution of the myelin compartment signal which is neglected in M2 but non-negligible in the short TE regime. For *in vivo* application of M2, new *in vivo* meGRE protocols need to be developed that allow for a longer maximal TE (e.g. ≥ 54 ms), where sufficient data points with dominant extra-cellular signal compartment are collected. However, estimating robust *in vivo* R_2,iso_* maps at these large TE_max_ values requires the correction of motion artefacts (Magerkurth et al., 2011), which is an interesting future project for itself.

Interestingly, the biggest discrepancy between *in silico* and *ex vivo* results for *β*_1_ was seen for the smallest maximal TE value at 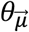 smaller than the magic angle (55°, Figure 10B). This is because *β*_1_ and *α*_1_ of the measured ex vivo data showed in this 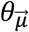 range an atypical 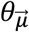 dependence: they decreased as a function of increasing 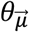 up to the magic angle. One might suspect that this atypical 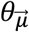 dependence of *β*_1_ has been artificially introduced by the higher-order model M2. However, the fact that it was also found for *α*_1_ makes it more likely that we have found a new orientation dependence of R_2_* that cannot be explained by the HCFM. A mechanism that could explain a reduction in R_2_* at the magic angle would be shortening of R_2_ due to the Magic Angle Effect in highly structured molecules like myelin sheaths (see (Bydder et al., 2007)). Since this phenomenon would be superimposed on the orientation dependence of R_2_* being investigated here, it may be particularly evident when the latter effect is negligible, i.e. at low 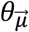.

### Considerations

Two of our findings might appear contradictory, at first glance: On the one hand, M2 can effectively capture the orientation independent component of R_2_* (i.e. estimate R_2,iso_*) at long maximal TEs (∼ 54 ms), indicating that the myelin-water pool can be neglected in this TE range. On the other hand, M2 cannot predict the fitted *β*_1_ whereas a heuristic expression that incorporates the contribution of the myelin water in *β*_1_ substantially improves the prediction. This contradiction can be resolved when considering the time-dependency of the orientation dependent and independent parameters of M2, i.e. *β*_2_ and *β*_1_ respectively (Equation 1). The *β*_2_ parameter scales with the square of time and thus captures the logarithm of the signal-decay at the higher-TE time-points where the contribution of the myelin water is negligible, whereas *β*_1_ scales linearly with time and thus captures the logarithm of the signal-decay at the smaller-TE time-points where the contribution of the myelin water cannot be neglected. Consequently, M2 can effectively separate-out the orientation dependency of R_2_* but at the same time might fail to predict *β*_1_ accurately. Future studies should aim to find a better derivation of M2 from the HCFM that does not neglect the contribution of the myelin water. From the perspective of interpretation, our heuristic expression of *β*_1_ might be particularly helpful because we found that it is a good proxy for the fitted *β*_1_.

To generate the *in silico* data, we employed simplifications to the HCFM when extended to multiple cylinders contained in an MRI volume with varying degree of dispersion. We assumed that the signal coming from multiple dispersed hollow cylinders is a super-position of the complex signal of multiple single hollow cylinders with a specific orientation to an arbitrary main orientation of the fibres. As a result, the near-field interaction of the cylinders was neglected. Moreover, the dephasing due to the myelin compartment was also assumed to be negligible (i.e., D_M_ ≈ 0). Nevertheless, the *in silico* data described the 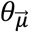 dependence of *α*_1_ and *β*_1_ as in the *ex vivo* data. This is seen across all dispersion regimes when using the long maximal TE protocol. As compared to previous studies where the dephasing process was more faithfully described in two dimensions (Hédouin et al., 2021; Xu et al., 2018), our model allowed for better control over the fibre dispersion in three dimensions via the Watson distribution parameter κ. Future work should investigate whether the validity of the in silico data could be improved by combining the approach of (Hédouin et al., 2021) in a three-dimensional simulation environment where the degree of fibre dispersion can be changed as well.

The *ex vivo* data possess two issues warranting discussion: (i) the coregistration of the diffusion results from the NODDI model into the GRE space and (ii) the use of the Watson dispersion from the NODDI model as a descriptor of the different fibre arrangements in the brain. Regarding the coregistration of the diffusion results from NODDI (see section 3.1.4), image interpolation could introduce a bias on the κ and, especially, 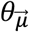 maps. The latter can be more affected in areas with strong angular gradients (e.g. a 0°-90° between two adjacent voxels), resulting in local over- or under-estimation of the 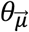 values. One solution is to ensure the acquisition of both MRI techniques (R_2_*w GRE and dMRI) occur with the specimen in the same MR system, with identical positioning, field-of-view and image resolution. For our study, this was not possible because we used a preclinical MR system to acquire the high-resolution diffusion MRI data whereas the meGRE data were acquired on a human 7 T system. Regarding the use of Watson dispersion from the NODDI model, this distribution cannot describe all existing fibre arrangements in the brain accurately, e.g., the crossing fibre arrangement. In the optic chiasm specimen such arrangements were only found in a few regions, e.g. at the crossing of the optical tract and optic nerve. Therefore, the contribution of such crossing-fibre voxels with estimated κ values in the range of highly to mildly dispersed fibres will be averaged-out with the single-fibre orientation voxels with similar κ values during the irregular binning pre-processing (section 3.3.1). However, this could result in an increasing standard deviation in the estimated *α*-parameters in the log-linear model and *β*-parameters in the log-quadratic model.

## 6. Conclusion

We showed that our recently introduced biophysical log-quadratic model of the multi-echo gradient-recall echo (meGRE) signal can effectively estimate the fibre-angular-orientation independent part of R_2_* (R_2,iso_*) for varying g-ratio values and fibre dispersions. Thus, it provides an attractive alternative to standard methods for deciphering the orientation-dependence of R_2_* that requires multiple acquisition with distinct positioning of the sample in the head-coil. Doing so would provide a more robust marker for neuroscientific studies in a broadly accessible manner. We also showed that the estimated linear time-dependent parameter of M2, *β*_1_, can be used to estimate the myelin water fraction (MWF) and g-ratio using a newly proposed heuristic expression relating *β*_1_ to microstructural tissue parameters including the myelin water signal. Importantly, we found that the proxy of R_2,iso_* cannot be estimated effectively with the log-quadratic model at lower echo time ranges (i.e. at maximal echo times smaller than 36 ms) that are typically used for whole-brain *in vivo* meGRE experiments. To make M2 usable for *in vivo* applications, future studies need to develop new meGRE protocols with longer TE_max_ (54 ms) that remain time efficient and motion robust. Finally, at echo time ranges smaller than 18 ms, an unexpected R_2_* orientation-dependence was found in the ex vivo dataset at angles below the magic angle: a decrease of R_2_* for increasing angles. Our HCFM based simulations were not able to model this angular dependence, which points towards a distinct mechanism in white matter that cannot be explained by the HCFM.

## Glossary

### A. Acronyms

#### a. Biophysical terms and model acronyms

AWF: (Intra-)Axonal water fraction
EWF: Extra-axonal water fraction
FVF: Fibre volume fraction
HCFM: Hollow cylinder fibre model
ICVF: Intra-cellular volume fraction (from NODDI)
MWF: Myelin water fraction

#### b. Magnetic resonance imaging and sequence acronyms

dMRI: Diffusion-weighted Magnetic Resonance Imaging
DWI: Diffusion-weighting Imaging
GRE: Gradient-recalled echo
meGRE: Multi-echo gradient-recalled echo
OC: Optic chiasm
R_2_*: Effective transverse relaxation rate
R_2,iso_*: Orientation independent or isotropic part of R_2_*
TE: Echo time
TE_max_: Maximal echo time

#### c. Hollow cylinder fibre model relevant acronyms

S_A_: Signal of the intra-axonal compartment
S_E_: Signal of the extra-axonal compartment
S_M_: Signal of the myelin compartment
S_N_: Sum of the signals of the non-myelinated (S_A_ and S_E_) compartments
S_C_: Sum of all the signal compartments (S_A_, S_E_ and S_M_)
R_2A_: Transverse relaxation rate of the intra-axonal compartment
R_2E_: Transverse relaxation rate of the extra-axonal compartment
R_2N_: Transverse relaxation rate of the non-myelinated compartments (R_2A_ = R_2E_ = R_2N_)
R_2M_: Transverse relaxation rate of the myelin compartment

### B. Symbols

#### a. In silico and ex vivo data

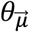: Angular orientation of the mean fibre bundle
*θ*_0_: First angular orientation or angular offset
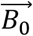: Main magnetic field
κ: Coefficient of dispersion (from Watson Distribution and NODDI)
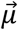: Vector of the mean fibre bundle
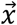: Vector of the individual cylinder in the simulated in silico data
*T*_*Diff,GRE*_: Transformation matrix from dMRI to GRE images
*T*_*GRE*:*i*,1_: Transformation matrix from GRE images at the i-th angular orientation measurement to the first angular orientation measurement

#### b. Model parameters and analysis

*α*_0_: Intercept parameter of M1
*α*_1_: Slope or linear parameter of M1
*β*_0_: Intercept of M2
*β*_1_: Slope of linear parameter of M2
*β*_1,nm_: *β*_1_ ground-truth value without myelin signal contribution (Equation 3)
*β*_1,m_: *β*_1_ ground-truth value with myelin signal contribution (Equation 4)
*β*_2_: Quadrature or second order parameter of M2
ε: Relative difference between fitted *β*_1_ and *β*_1,theory_ (Equation 9)
ε_m_: Relative difference between fitted *β*_1_ and predicted *β*_1,nm_
ε_nm_: Relative difference between fitted *β*_1_ and predicted *β*_1,m_
nRMSD: Normalised root-mean-squared deviation (Equation 7)
ΔRMSD: Normalised root-mean-squared deviation difference (Equation 8)
M1: Log-linear model (Equation 2)
M2: Log-quadratic model (Equation 1)

## 7. Acknowledgment

This work was supported by the German Research Foundation (DFG Priority Program 2041 “Computational Connectomics”, [AL 1156/2-1;GE 2967/1-1; MO 2397/5-1; MO 2249/3–1], by the Emmy Noether Stipend: MO 2397/4-1) and by the BMBF (01EW1711A and B) in the framework of ERA-NET NEURON and the Forschungszentrums Medizintechnik Hamburg (fmthh; grant 01fmthh2017). The research leading to these results has received funding from the European Research Council under the European Union’s Seventh Framework Programme (FP7/2007-2013) / ERC grant agreement n° 616905. MFC is supported by the MRC and Spinal Research Charity through the ERA-NET Neuron joint call (MR/R000050/1). The Wellcome Centre for Human Neuroimaging is supported by core funding from the Wellcome [203147/Z/16/Z]. The Max Planck Institute for Human Cognitive and Brain Sciences has an institutional research agreement with Siemens Healthcare. NW holds a patent on acquisition of MRI data during spoiler gradients (US 10,401,453 B2). NW was a speaker at an event organized by Siemens Healthcare and was reimbursed for the travel expenses.

## 9. Appendix

### 9.1 Hollow cylinder fibre model in detail

The hollow cylinder fibre model (HCFM) proposes an analytical approximation of the angular orientation 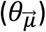 dependency of the GRE signal to 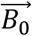. This approximation establishes that the total MR signal comes from water molecules in an *infinitely long* hollow cylinder affected by the diamagnetic myelin sheath (Liu, 2010). The diamagnetic myelin sheath perturbs magnetically the water molecules from three distinguishable compartments: (1) the intra-axonal (S_A_), (2) myelin (S_M_) and (3) extra-cellular (S_E_) compartments. Then, the total MR signal, S_C_, is defined as:

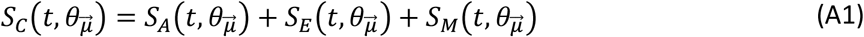

where the signal decay coming from each compartment, as defined in (Wharton and Bowtell, 2012) and (Wharton and Bowtell, 2013), are defined as a function of time (t) and 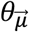:

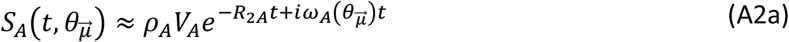

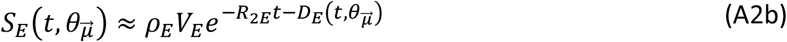

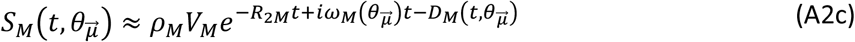

In these compartmental equations, ρ, R_2_ and V are respectively the proton density, transverse relaxation rate and volumes for each compartment (defined with the corresponding sub-indices). The functions ω_A_ and ω_M_ are the (local) frequency offset of the intra-axonal and myelin water molecules produced by the myelin susceptibility (from (Wharton and Bowtell, 2012) and (Duyn, 2014)), defined as:

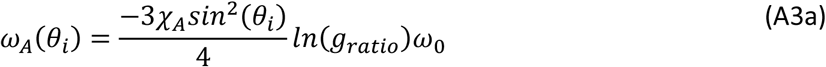

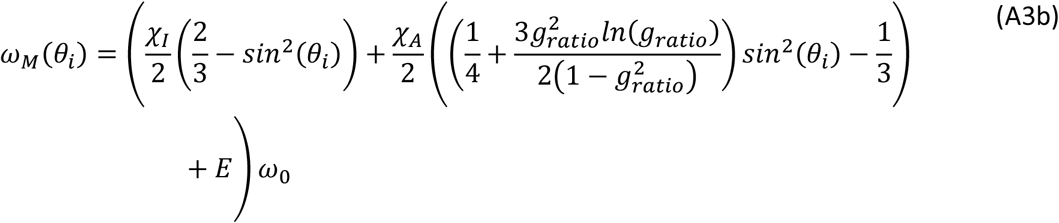

where χ_I_ and χ_A_ are the isotropic and anisotropic magnetic susceptibilities of the myelin sheath (in ppm), E is the exchange factor between compartments (in ppm) and ω_0_ is the Larmor frequency (= 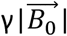, in MHz, with γ the gyromagnetic ratio) of the water molecules. The D_E_ and D_M_ functions are the dephasing in the extracellular and myelin compartments across the voxel. D_E_ is defined, in the work of (Wharton and Bowtell, 2013), as a piece-wise function using the approximation introduced by (Yablonskiy and Haacke, 1994) and discussed in section 9.2. The D_M_ function is neglected in this study.

### 9.2. Analytical expression of the dephasing component (D _E_) of the extracellular compartment (S_E_)

Yablonskiy and Haacke, (1994) proposed the analytical expression for the magnetic dephasing of a medium due to the presence of cylindrical dephasors (defined as cylinders with a different magnetic susceptibility than the medium) oriented with an angle 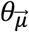 to 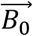 defined as:

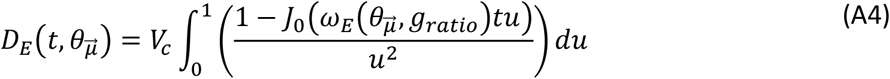

Where V_c_ is the cylinder’s volume (equal to the fibre volume fraction, FVF), J_0_ is the zeroth-order Bessel’s function of the First Kind, u is the variable of integration and ω_E_ is the frequency offset in the extracellular space. The latter is defined as:

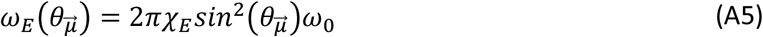

Where χ_E_ is the mean susceptibility of the myelin sheet, defined as (χ_I_ + 0.25 χ_A_)(1 – g^2^_ratio_). In their work, Equation A3 was approximated for two-time scales divided by the so-called critical time (α in (Wharton and Bowtell, 2013)), defined as:

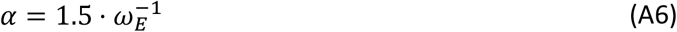

For times shorter than the critical time, the dephasing function is approximated by a quadratic function, while for times longer than the critical time this function becomes linear. The corresponding analytical expressions (Yablonskiy and Haacke, 1994) are:

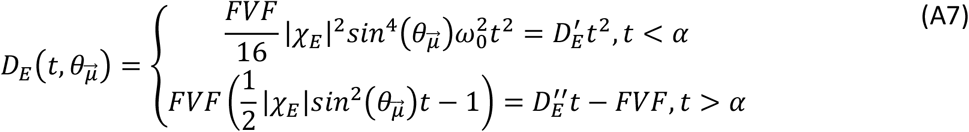

Where 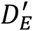 and 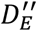 are expressions having all the parameters that are not time dependent, including 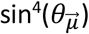 and 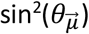, respectively. This simplified expression, especially the quadratic approximation, is used later (section 9.3). However, this piecewise approximation has a discontinuity at this critical time, as observed in Figure A1. To avoid this discontinuity when D_E_ overpasses the critical time for the *in silico* data, we used an analytical solution to Equation A4. This solution was performed in Mathematica 12 (Wolfram Research, Inc., Champaign, IL (2020)), giving the following expression:

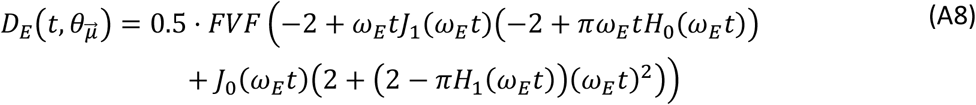

In where J_1_ is the first-order Bessel’s function of the First-Kind, and H_0_ and H_1_ are the zeroth and first-order Struve functions ((Struve, 1882) and (Aarts and Janssen, 2016)), respectively. The offset frequency in the extracellular space (ω_E_) is dependent on the mean angular orientation and the g-ratio, as defined in (Yablonskiy and Haacke, 1994) and (Wharton and Bowtell, 2012).

**Figure A 1:**
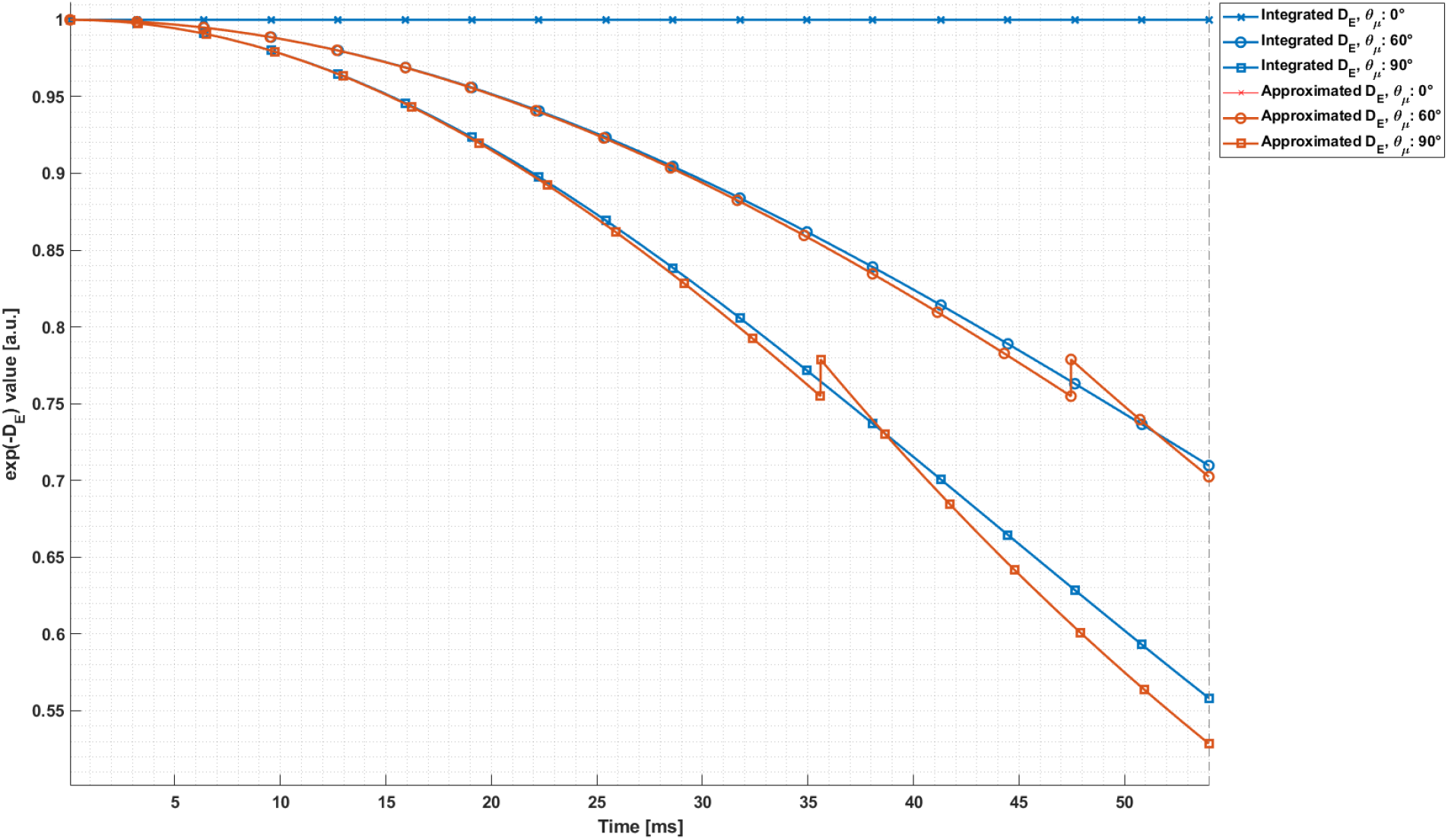
Signal decay in the extra-cellular compartment due only to dephasing (D_E_) using different D_E_ functions. The signal decay (i.e., exp(-D_E_) in Equation A2b) was evaluated in function of time (in ms) and at three different angular orientations (0°, 60° and 90°). Two expressions for the D_E_ function (Equation A4) were used: the analytical solution given in Equation A8 (Integrated D_E_, blue curve) and the piece-wise approximation proposed in the work of Yablonskiy et al. 1994 in Equation A7 (approximated D_E_, orange curve). Both functions were evaluated using the simulation values (section 9.4, Tables A1 to A3).

### 9.3. Analytical interpretation of the log-quadratic model (M2) and approximation with myelin compartment added

The log-quadratic model (M2) is derived from the signal equation from the HCFM with neglected myelin-water signal (S_M_, Equation A4). This signal is neglected due to short T_2_ and small volume of this compartment (see Wharton and Bowtell, 2013). The magnitude of the remaining signal of the non-myelin compartments (S_N_) is defined as:

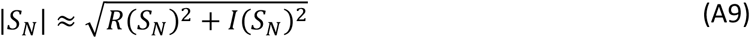

where *R* and *I* are the real and imaginary components of S_N_ and S_N_ is defined as follows:

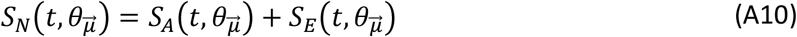

Evaluating Equation A10 with Equations A2a-b resulted in:

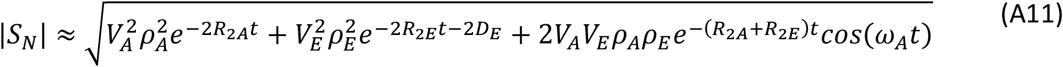

Using the natural logarithm function (ln(x)) of the above equation results in:

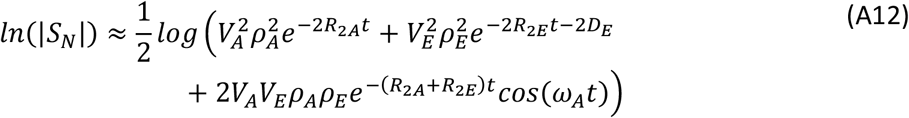

This expression can be linearised if the three functions related to time, i.e. the transverse relaxation rates (e.g. 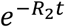), the frequency offset of the intra-axonal compartment (cos(ω_A_t)) and dephasing of the extra-axonal compartment (D_E_), are *sufficiently small* to be approximated using the 1^st^ and 2^nd^ order of the Taylor expansion, respectively, as follows:

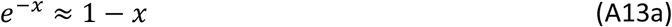

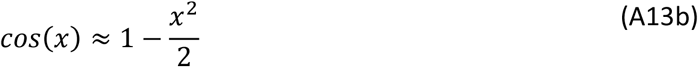

If these conditions are fulfilled, the logarithm function of Equation A12 can be approximated by a 2^nd^ order Taylor expansion in time, resulting in:

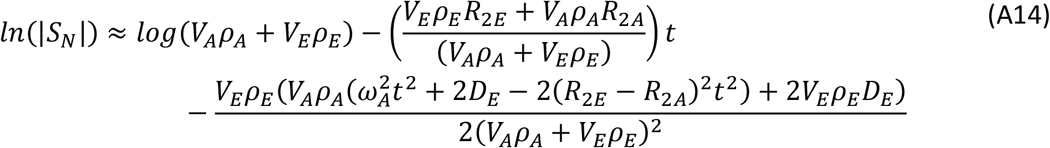

If the quadratic approximation for D_E_ is used (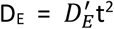, Equation A7), this expression can be summarized as:

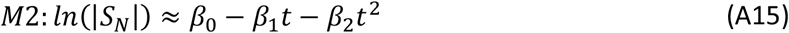

Where:

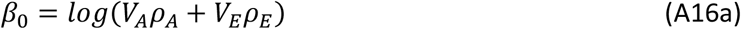

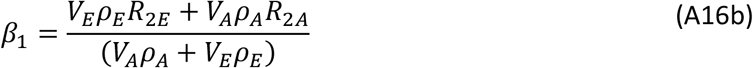

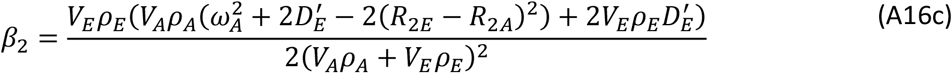

In the scenario where R_2A_ is equal to R_2E_, the analytical expression for *β*_1_ becomes *β*_1,*nm*_ (Equation 3, section 2.2).

The proposed heuristic analytical expression of *β*_1_ in Equation 4, *β*_1,*m*_, was motivated by taking Equation A16b and incorporating the myelin compartment information (V_M_, ρ_M_ and R_2M_) in a similar manner, resulting in the following expression:

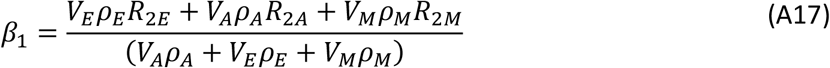

This expression can also be derived as the linear component in time by keeping the contribution of the myelin compartment in Equation A10, i.e. using S_C_ from Equation A9 instead of S_N_, and performing a Taylor expansion in time. While *β*_1_in Equation A17 (or *β*_1,*m*_ in Equation 4) turned out to explain better the *in silico* fitted *β*_1_ than the *β*_1,*nm*_ (Equation 3 and A16b, see Figure 7), the validity range of the second-order approximation of the entire signal S_C_ with the added myelin compartment is highly restrictive as a function of time and cannot be used for the experimental parameters used here (data not shown).

Equation A17 can be re-written as a function of the myelin water fraction (MWF), axonal water fraction (AWF) and extra-axonal water fraction (EWF), defined as:

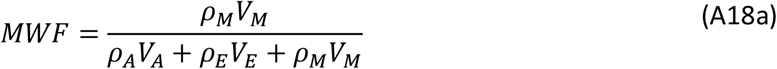

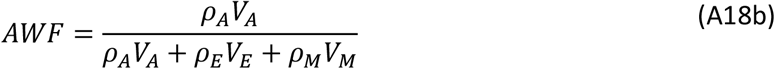

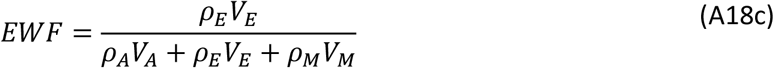

Since the sum of the water fractions are equal to 1, *β*_1_ becomes:

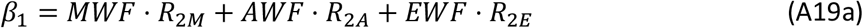

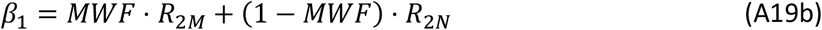

Where Equation 4 (or A19b) is obtained if we assume in Equation A19a that the relaxation rate in the intra- and extra-cellular water is the same: *R*_2*A*_ = *R*_2*E*_ = *R*_2*N*_.

### 9.4. *In silico* data setup: simulation parameters, SNR and anisotropic binning

The *in silico* MR data was simulated using the HCFM for each hollow cylinder. The fixed parameters were obtained from (Wharton and Bowtell, 2013) and listed as follows:

**Table A 1:** Fixed microstructural parameters used to create the in silico data from (Wharton and Bowtell, 2013) in section 3.2. *Proton densities were scaled by a factor of 5000 but they kept the same proton density proportion between the non-myelinated and myelinated compartments (1:0.7).

| Parameter | Value | Parameter | Value |
| --- | --- | --- | --- |
| Anisotropic and isotropic susceptibilities ($\chi_A$ and $\chi_I$) | -0.1 ppm | Larmor frequency at 7 T ($\omega_0$) | $1.873 \cdot 10^6$ rad/ms |
| Exchange (E) | 0.02 ppm | Fibre volume fraction (FVF) | 0.5 n. u. |
| Proton density intra- and extra- axonal compartments ($\rho_A$ and $\rho_E$)* | 5000 a. u. | Proton density myelin compartment ($\rho_M$)* | 3500 a. u. |

Other fixed parameters were obtained from (Dula et al., 2010b) and they are listed as follows:

**Table A 2:** Fixed microstructural parameters used to create the in silico data (section 3.2) obtained from (Dula et al., 2010a).

| Parameter | Value | Parameter | Value |
| --- | --- | --- | --- |
| $R_2$ intra- and extra- axonal compartments ($R_{2A} = R_{2E} = R_{2N}$) | $18.53 \text{ s}^{-1}$ | $R_2$ myelin compartment ($R_{2M}$) | $75.41 \text{ s}^{-1}$ |

The variable parameters, or parameter space, of the *in silico* MR data are listed as follows:

**Table A 3:** Variable microstructural and physical parameters, or parameter space, used to create the in silico data (section 3.2). The two extreme values for g-ratio, 0.66 and 0.8, were found in (Emmenegger et al., 2021) and (Wharton and Bowtell, 2013), respectively. The mean value of 0.73 was arbitrarily defined.

| Parameter | Value | Parameter | Value |
| --- | --- | --- | --- |
| Angular orientation ($\theta_{\vec{\mu}}$) | 2°:2°:90° | g-ratio | 0.66, 0.73, 0.8 |
| Index of fibre dispersion ($\kappa$) | 0.001:0.1:6.0 | Time | 3.25:3.25:53.5 ms |

To make the *in silico* as-similar-as possible to the *ex vivo* data, noise was added to the signal decay of the *in silico* data, in such a way that the *in silico* SNR is like the SNR seen in the *ex vivo* GRE data. For that, the *ex vivo* SNR was calculated by dividing the signal of the white matter region of the OC and the standard deviation of the background in (image) magnitude space. No noise correlation correction was performed in this calculation given the coil having 2 receiver channels. As a result, an average SNR value of 112 was obtained for this region (section 3.2), and with this SNR value, a complex random Gaussian noise was added to the *in silico* data as follows:

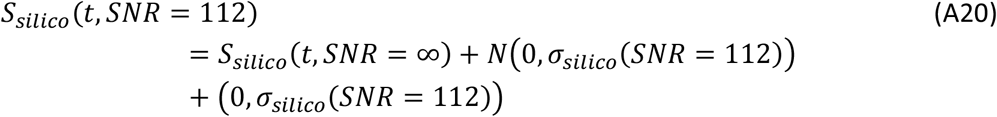

Where N(0,σ) is the Normal distribution with mean 0 and the standard deviation defined by:

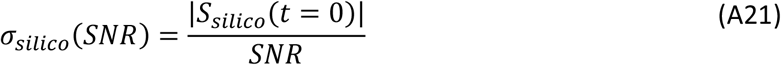

Where the magnitude signal is divided by the desired SNR at time 0 (|S_silico_(t = 0)|).

With the noise added, the magnitude of the *in silico* MR signal at SNR = 112 was obtained:

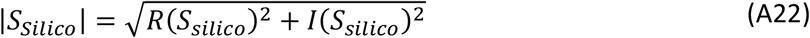

To compare the *in silico* data analysis across the 5000 signal decays per simulated g-ratio, sampled κ and 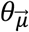 to the irregularly binned *ex vivo* data analysis (section 3.3.1 and Figure 5B), the *α*-parameters and *β*-parameters from the *in silico* data required three consecutive averaging-steps: (1) an averaging across the 5000 samples, resulting in the sampled-averaged 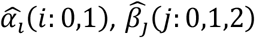 and their standard deviations *sd*(*α*_*i*_), *sd*(*β*_*j*_) per sampled κ value and 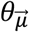. (2) A weighted averaging across κ values per each 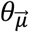 irregular bin of the *ex vivo* data in each κ range. For that, it was obtained the distribution of the κ values from the voxels contained in each of the 20 defined 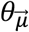 irregular bins. The 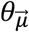 range per bin and κ range is given in Table A.4. Then, all the obtained distributions were averaged per κ range (Figure A2 from A to C) to remove possible influence of the irregular 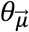 bins on κ. The standard deviation from this average was calculated, normalised and used later (referred as the sd(*P*(*κ*_*l*_)) in Equation A28). Next, a probability distribution, *P*(*κ*_*l*_), was fitted accordingly (Figure A2 from D to F) and the weighted averaging on 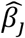 (the same procedure is performed for 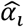) was calculated as follows:

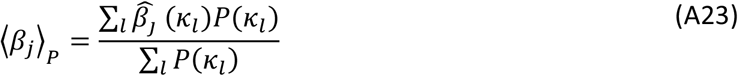

Where the expression for *P*(*κ*_*l*_) was heuristically chosen and varied per each fibre dispersion (κ range): a Beta distribution for the highly dispersed fibres (κ < 1 range, Figure A2-D), defined as:

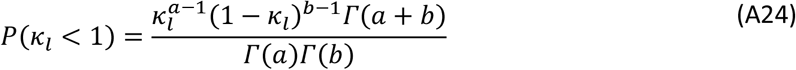

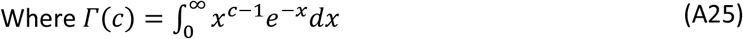

Is the Gamma function. The coefficients a and b estimated for this range were 3.145 and 1.234, respectively. Given the clear half-shaped normal distribution, a Half-Normal distribution for the mildly dispersed fibres (1 ≤ κ < 2.5 range, Figure A2-E) was used, defined as:

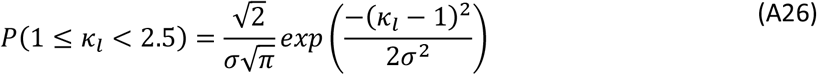

The coefficients μ and σ were 0 and 0.4498, respectively. And given the fast decay of the values at the beginning of the distribution, an Exponential distribution for the highly aligned fibres (2.5 ≤ κ range, Figure A2-F) was used, defined as:

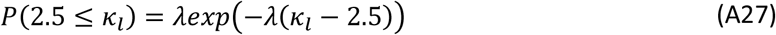

The coefficient λ was 0.2241. The standard deviation of ⟨*β*_*j*_ ⟩_*P*_ was also estimated by error-propagating the *sd*(*β*_*j*_) weighted by *P*(*κ*_*l*_) and its standard deviation sd(*P*(*κ*_*l*_)), as follows:

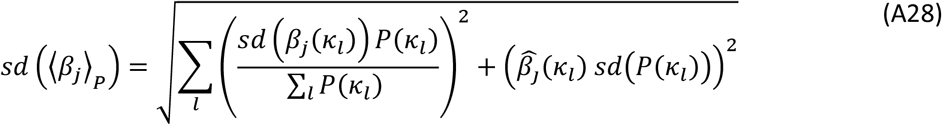

While the first squared term requires the normalisation factor (∑_*l*_ *P*(*κ*_*l*_)) because the weights *P*(*κ*_*l*_) are not normalised, the second is not needed since sd(*P*(*κ*_*l*_)) is already normalised. Finally, the ⟨*β*_*j*_⟩_*P*_ and *sd* (⟨*β*_*j*_⟩_*P*_), and the ⟨*α*_*i*_⟩_*P*_ and *sd*(⟨*α*_*i*_⟩_*P*_) (as in Equation A28) were averaged and error-propagated, respectively, as a function of the 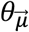 values for each irregular bin.

**Figure A 2:**
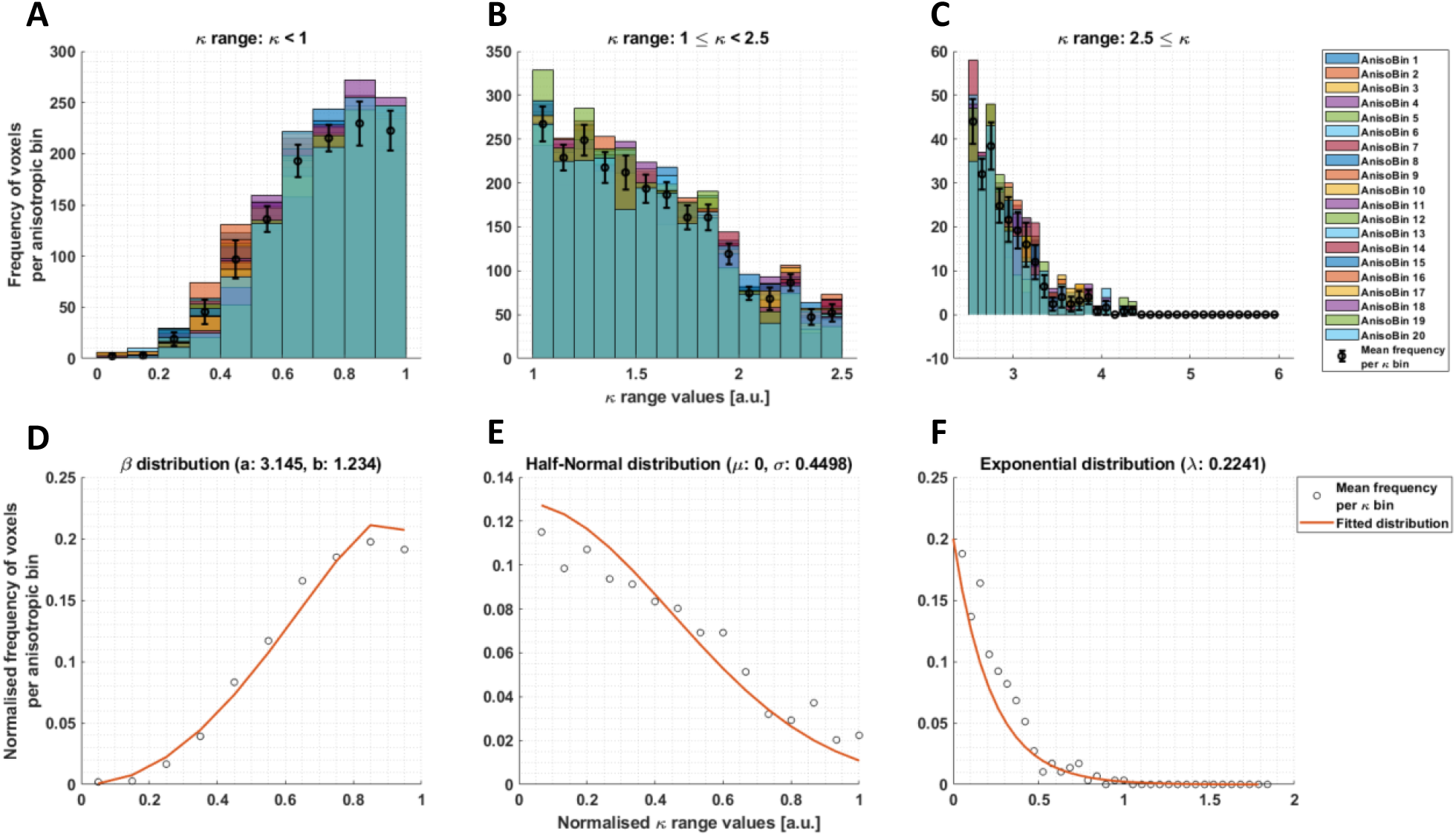
Assembling the in silico data across the simulated κ ranges and angular 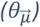 anisotropic bins. To make the in silico data comparable to the ex vivo data, the frequency of voxels as a function of κ was obtained per defined 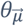 irregular bin in the ex vivo data (Figure 3). This was performed for the 20 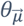 bins (AnisoBin X, with X the corresponding bin from 1 to 20, see Table A.4) and per fibre dispersion (κ range): highly dispersed (κ < 1, A), mildly dispersed (1 ≤ κ < 2.5, B) and negligibly dispersed (κ ≥ 2.5, C) fibres. The mean and standard deviation across histograms were obtained (error bars). The means were normalised with respect to the cumulated value (i.e., sum of all the mean values) and fitted with a continuous function (P(κ_l_)) per κ range, previously normalised: a beta distribution for κ < 1 (D), half-normal distribution for 1 ≤ κ < 2.5 (E) and exponential distribution for κ ≥ 2.5 (F). The standard deviation was also normalised by the cumulated value per κ range and used as the standard deviation of the continuous distributions (sd(P(κ_l_))).

**Table A 4:** Range of angles 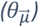 defined by [min, max] values, contained in each 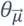 irregular bin per fibre dispersion (κ range) in the ex vivo data (section 3.3.1). The angular offset, θ_0_ (see section 3.3.1), is defined as the angular average of the 1^st^ irregular bin, resulting in 17.3° (κ < 1), 20.4° (1 ≤ κ < 2.5) and 22.9° (2.5 ≤ κ).

| Dispersion range<br>Irregular bin | $1 < \kappa$ | $1 \leq \kappa < 2.5$ | $2.5 \leq \kappa$ |
| --- | --- | --- | --- |
| #1 and ($\theta_0$) | [0, 29.3]$^\circ$<br>(17.3$^\circ$) | [0, 26.5]$^\circ$<br>(20.4$^\circ$) | [0, 30.7]$^\circ$<br>(22.9$^\circ$) |
| #2 | [29.3, 39.3]$^\circ$ | [26.6, 36.8]$^\circ$ | [30.8, 40.8]$^\circ$ |
| #3 | [39.4, 45.9]$^\circ$ | [36.9, 43.5]$^\circ$ | [40.9, 47.2]$^\circ$ |
| #4 | [45.6, 50.8]$^\circ$ | [43.6, 48.6]$^\circ$ | [47.3, 52.1]$^\circ$ |
| #5 | [50.9, 55.0]$^\circ$ | [48.7, 52.9]$^\circ$ | [52.2, 56.2]$^\circ$ |
| #6 | [55.1, 58.6]$^\circ$ | [53.0, 56.6]$^\circ$ | [56.3, 59.7]$^\circ$ |
| #7 | [58.7, 61.8]$^\circ$ | [56.7, 60.0]$^\circ$ | [59.8, 62.7]$^\circ$ |
| #8 | [61.9, 64.7]$^\circ$ | [60.1, 63.0]$^\circ$ | [62.8, 65.6]$^\circ$ |
| #9 | [64.8, 67.4]$^\circ$ | [63.1, 65.9]$^\circ$ | [65.7, 68.3]$^\circ$ |
| #10 | [67.5, 70.0]$^\circ$ | [66.0, 68.5]$^\circ$ | [68.4, 70.7]$^\circ$ |
| #11 | [70.1, 72.3]$^\circ$ | [68.6, 71.0]$^\circ$ | [70.8, 72.9]$^\circ$ |
| #12 | [72.4, 74.6]$^\circ$ | [71.1, 73.5]$^\circ$ | [73.0, 75.2]$^\circ$ |
| #13 | [74.7, 76.7]$^\circ$ | [73.6, 75.7]$^\circ$ | [75.3, 77.2]$^\circ$ |
| #14 | [76.8, 78.7]$^\circ$ | [75.6, 77.8]$^\circ$ | [77.3, 79.2]$^\circ$ |
| #15 | [78.8, 80.6]$^\circ$ | [77.9, 79.8]$^\circ$ | [79.3, 81.1]$^\circ$ |
| #16 | [80.7, 82.4]$^\circ$ | [79.9, 81.7]$^\circ$ | [81.2, 82.8]$^\circ$ |
| #17 | [82.5, 84.1]° | [81.8, 83.6]° | [82.9, 84.4]° |
| #18 | [84.2, 85.7]° | [83.7, 85.3]° | [84.5, 85.9]° |
| #19 | [85.8, 87.3]° | [85.4, 87.3]° | [86.0, 87.4]° |
| #20 | [87.4, 90]° | [87.4, 90]° | [87.5, 90] |

