## Supplementary Figure 1 for "Deciphering the fibre-orientation independent component of R_2_* (R_2,iso_*) in the human brain with a single multi-echo gradient-recalled-echo measurement under varying microstructural conditions"

### Supplementary material

#### 1. Supplementary figure

Figure S1 shows eight of the sixteen angular-measured optic chiasm maps after coregistration (section 3.1.4 in main manuscript). For each measurement, the estimated voxel-wise angular ($\theta_{\vec{\mu}}$) map with the corresponding angular orientation of the external magnetic field ($\theta_{\overrightarrow{B_0}}$) are shown. The $\theta_{\overrightarrow{B_0}}$ was estimated as $\theta_{\overrightarrow{B_0}} = acos\left(\overrightarrow{B_0}(\theta_i) \cdot \overrightarrow{B_0}(\theta_0)\right)$. Middle and bottom rows also show the estimated $R_2$* via $\alpha_1$ in M1 (Equation 2) and $R_{2,iso}$* via $\beta_1$ in M2 (Equation 1).

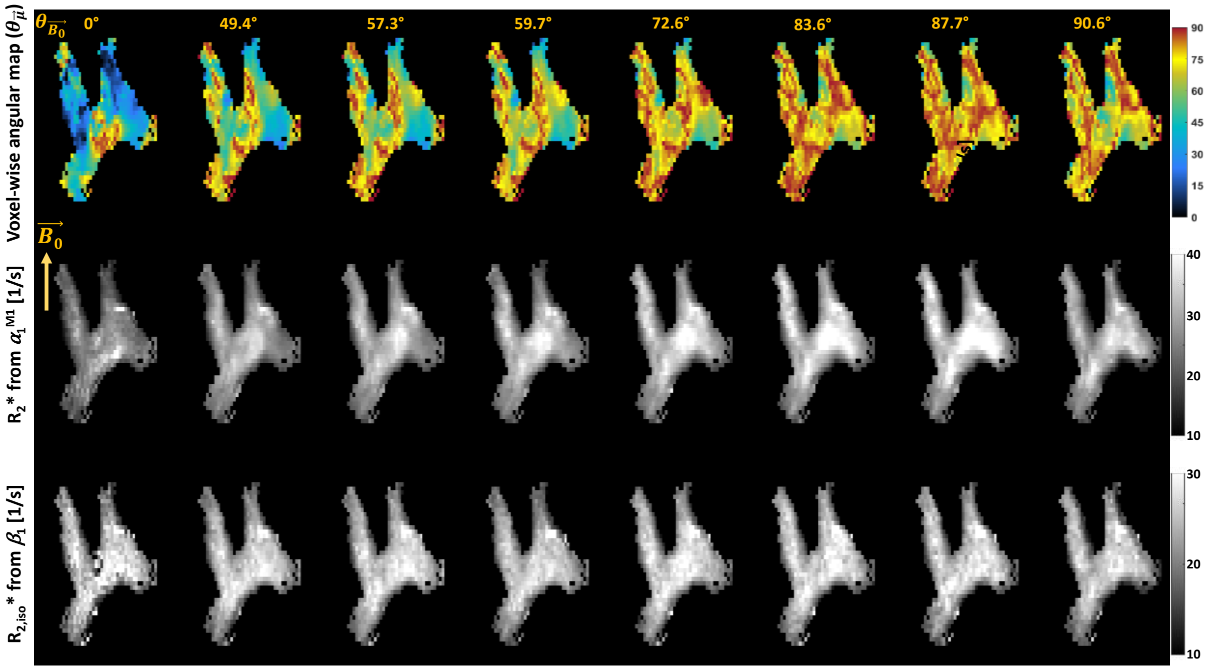


*Figure S 1: $R_2$* angular ($\theta_{\vec{\mu}}$) dependency in a coronal section of the ex vivo specimen from 8 of the 16 angular measurements. They are sorted using the calculated $\overrightarrow{B_0}(\theta_i)$ angular elevation ($\theta_{B_0}$ in the inset) after coregistration: The first row shows the voxel-wise angular $\theta_{\vec{\mu}}$ map constrained between 0° and 90°. Second row is the estimated $R_2$* via $\alpha_1$ parameter maps analysed with M1 (Equation 2). Third row shows the estimated $R_{2,iso}$* via $\beta_1$ parameter maps analysed with M2 (Equation 1).*
